# Decision-related signals in the presence of nonzero signal stimuli, internal bias, and feedback

**DOI:** 10.1101/118398

**Authors:** Daniel Chicharro, Stefano Panzeri, Ralf M. Haefner

## Abstract

Understanding the nature of decision-related signals in sensory neurons promises to give insights into their role in perceptual decision-making. Those signals, traditionally quantified as choice probabilities (CP), are well-understood in a feedforward framework assuming zero-signal trials with no choice bias. Here, we extend this understanding by analytically solving models of choice-related signals that account for informative stimuli, choice bias, and importantly, feedback signals reflecting either internal states, such as attention or belief, or the outcome of the decision process. First, we relate CPs to Choice Triggered Averages (CTAs), which quantify choice-related average changes in neural responses, and show that both have general expressions valid for activity-choice covariations of both feedforward or feedback origin. These expressions allow a meaningful calculation of CPs across all trials, including non-zero signal trials. Second, we derive how CPs and CTAs depend on feedforward and feedback weights and on noise correlations under several plausible model architectures. Third, we examine different types of feedback signals, related to predictive coding, probabilistic inference, and attention, and we predict how CPs and CTAs depend in each case on the stimulus signal level and on the neural tuning properties. Finally, we show that measuring both CPs and CTAs offers complementary information about the origin of choice-related signals, especially when studying temporal changes of activity-choice covariations across the trial time. Overall, our work provides new analytical tools to better understand the link between sensory representations and perceptual decisions.

## 1 Significance Statement

The covariation between responses of sensory neurons and behavioral choices has traditionally been studied using stimuli uninformative about the correct decision, with the aim to separate the choice-related component of the responses from stimulus-driven activity. Furthermore, although recent experimental findings suggest an important contribution of feedback signals to choice-related activity, theoretical models providing analytical solutions of the measures quantifying this activity have focused on describing the feedforward influence that trial-to-trial noise variability in the sensory responses has in the decision. Our work covers this gap presenting and analyzing general analytical solutions which are valid in the presence of informative stimuli, choice bias, and feedback signals, thus providing new tools to interpret choice-activity covariations and understand their origin and functional role.

## 2 Introduction

Understanding how responses of sensory neurons contribute to perceptual decisions is a fundamental question in systems neuroscience. Much research addressing this question is based on examining the covariation between the responses of single neurons and behavioral choices. In the most established feedforward model (Shadlen et al., 1996; Gold and Shadlen, 2007), these activity-choice covariations have been interpreted based on the feedforward influence that trial-to-trial noise variability in sensory responses has on the decision. Within this feedforward framework, model simulations (Shadlen et al., 1996; Cohen and Newsome, 2009; Nienborg and Cumming, 2010) and analytical calculations (Haefner et al., 2013) have explained the activity-choice covariations of single neurons in terms of the combination of the distribution of feedforward readout weights and the structure of noise correlations across the neuronal population.

However, whether activity-choice covariations are due to causal influences of sensory neurons responses on the decision via feedforward pathways, or due to decision-related feedback signals influencing sensory responses, is a much-debated question (Cumming and Nienborg, 2016). Experimental (Nienborg and Cumming, 2009) and modeling (Wimmer et al., 2015; Haefner et al., 2016) evidence supports that feedforward and feedback signals can contribute differently to activity-choice covariations during different times of the decision process, and top-down modulations of sensory responses play an important role in a variety of theories of sensory coding, such as predictive (Rao and Ballard, 1999) or probabilistic (Lee and Mumford, 2003; Haefner et al., 2016; Tajima et al., 2016) coding.

In the context of two-choice discrimination tasks, choice probabilities (CP) have been the most prominent measure to study and quantify activity-choice covariations (Britten et al., 1996; Parker and Newsome, 1998; Nienborg et al., 2012). To separate the component of neural responses related to choice from stimulus-driven activity, CPs have been either calculated from zero-signal stimuli trials or, when using trials in which an informative stimulus is presented, discounting the estimated stimulus-driven response component (Kang and Maunsell, 2012). However, how the interplay of these two components that determine the choice affects CPs has not been examined. Thus, a better understanding of how the interaction of stimulus-driven activity and internal responses variability influences activity-choice covariations, as well as of the contribution of both feedforward and feedback signals, requires significantly extending the mathematical framework that so far has been used to interpret CPs. In particular, closed-form solutions for CPs (Haefner et al., 2013) only exist for very specific assumptions: a feedforward model, for trials with an equal number of choices for either of the two choices, that is, for unbiased decisions in the presence of zero-signal stimuli.

Here, we significantly extend the approach of Haefner et al. (2013) by deriving analytical relationships accounting for informative non-zero signal stimuli, choice bias, and importantly, feedback signals, either from generic internal states or specifically associated with the decision process. We introduce analytical expressions that can be used to characterize activity-choice covariations regardless of their origin, either due to feedforward or feedback pathways. In particular, we analytically relate CPs to other measures of covariations including Choice Correlation (CC) (Pitkow et al., 2015) and Choice Triggered Average (CTA) (Haefner, 2015). Furthermore, our analytical results also apply to detect probability, equivalent to CP but for detection tasks (Bosking and Maunsell, 2011; Smolyanskaya et al., 2015; Nienborg et al., 2012).

This generalized analytical framework contributes to our understanding of the origin of activity-choice covariations in several ways. First, it shows how to compute CPs across trials with different signal strength. Second, for simple models comprising feedforward and feedback signals, we derive how CPs and CTAs depend on feedforward and feedback weights as well as on noise correlations. Third, we discuss how to combine different measures to characterize the changes in activity-choice covariations. Fourth, we illustrate the signatures of different types of feedback with different functional roles, such as feedback related to predictive coding, probabilistic inference, or attentional modulations. Overall, we provide novel analytical tools to better study activity-choice covariations, moving from the quantification of single CP values to studying covariations as a function of stimulus signal levels and the cell’s tuning properties, deriving CP and CTA expressions that can accommodate different assumptions about whether the covariations originate from feedforward signals, feedback signals, or both.

## 3 Methods

This section contains a more detailed description of the derivations followed to obtain analytical solutions for the activity-choice covariation measures and a description of generative models representative of different sources of covariations. To make the paper as accessible as possible, the Results section is self-contained. Readers not interested in the detailed derivations can skip ahead directly to Results.

### 3.1 An exact CP solution for the threshold model

We here derive an analytical CP expression valid in the presence of informative stimuli, of decision-related feedback, and of top-down sources of activity-choice covariation, such as prior bias, trial-to-trial memory, or internal state fluctuations. We follow Haefner et al. (2013) and assume a threshold model of decision making, in which the choice *D* is triggered by comparing a decision variable *d* with a threshold *θ*, so that if *d* > *θ* choice *D* =1 is made, and *D* = − 1 otherwise. To obtain an exact solution of the CP we assume that the distribution *p*(*r_i_, d*) of the responses *r_i_* of cell *i* and the decision variable *d* can be well approximated by a bivariate Gaussian. Given these assumptions we calculate the choice probability, defined as

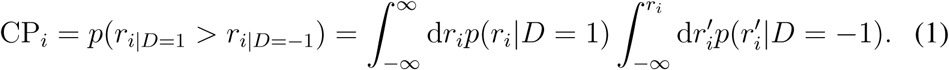

The CP is a quantification of the difference between the distributions of the responses for each choice, *p*(*r_i_*|*D* = −1) and *p*(*r_i_*|*D* = 1). It can be interpreted as the probability of being correct for an ideal observer that assigns *D* = 1 to the sample with higher value from two samples drawn together, one from each of the two distributions. If there is no dependence between the choice and the responses this probability is CP = 0.5. Following Haefner et al. (2013) (Supplementary Material *S*1) we get the conditional distribution

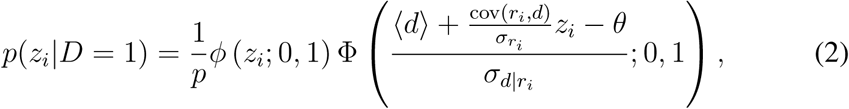

where a more parsimonious expression is obtained using the z-score 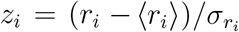 This distribution is a skew-normal (Azzalini, 1985), where *ϕ*(⋅; 0,1) is the standard normal distribution with zero mean and unit variance, and Φ(⋅; 0,1) is its cumulative function. Furthermore, cov(*r_i_*, *d*) is the covariance of *r_i_* and *d*, *σ_d_|r_i_* is the conditional standard deviation of *d* given *r_i_*, and the probability of *D* = 1 is

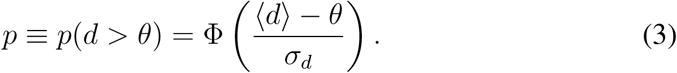

We will refer to *p* as the *Choice Ratio.* Intuitively, *p* increases when the mean of the decision variable 〈*d*〉 is higher than the threshold *θ*, and decreases when its standard deviation *σ_d_* increases. Consistently, for an uninformative stimulus, *p* = 0.5. Eq.2 can be synthesized in terms of *p* and the correlation coefficient *ρ_r_i__d*, which was named by Pitkow et al. (2015)*Choice Correlation* (CC). In particular, defining 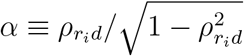 and 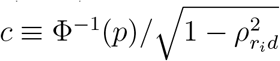 we get

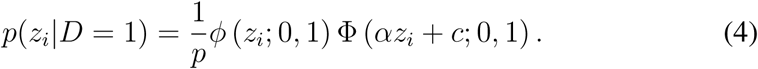

The CP is completely determined by *P*(*z_i_*|*D* = –1) and *P*(*z_i_*|*D* = 1), and since these distributions depend only on *p* and on the correlation coefficient *ρ_r_i__d*, the CP itself is a function of only these two quantities. Plugging the distribution of Eq.4 into the definition of the CP (Eq.1) we get

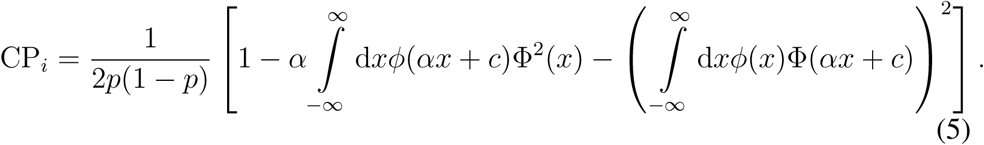

This expression is derived analogously to Eq. *S* 1.2 in Haefner et al. (2013), and generalizes the case examined there, which corresponds to *c* = 0. We now need some results involving integrals of normal distributions:

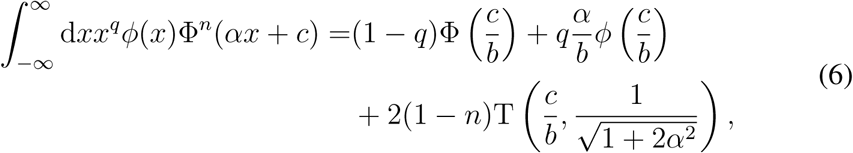

where 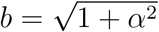 Here T is the Owen’s T function introduced by Owen (1956) and the equality above is valid for the cases *q =* 0, *n* =1,2, and *q =* 1 *,n =* 1 used by us. Using the equality for *q* = 0, *n* = 1,2 into Eq.5 leads to

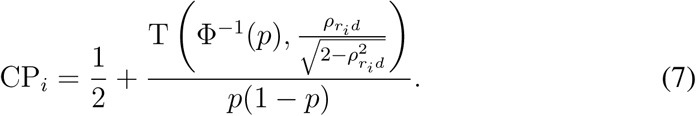

For an uninformative stimulus (*p* = 0.5), the function T reduces to the arctangent and the exact result obtained in Haefner et al. (2013) is recovered. The dependence on *ρ_r_i__d* can be intuitively understood because under the Gaussian assumption the covariance reflects all the dependence between the responses and the decision variable *d*. The dependence on the choice ratio reflects the influence of the threshold mechanism, which maps the dependence of *r_i_* with *d* into a dependence with choice *D* by partitioning the space of *d* in two regions. We will study this exact solution but, like Haefner et al. (2013), we mostly focus on a linear approximation derived in the limit of a small *ρ_r_i__d*. The rationale for this approximation is that experimentally estimated CPs are usually close to 0.5 (e. g.Britten et al., 1996; Nienborg et al., 2012), indicating that *ρ_r_i__d* is small, but we will see that the approximation is robust for a wide range of *ρ_r_i__d* values. For simplicity, we here present only a restricted derivation of this approximation that follows directly from the exact solution above. However, the same approximation could be derived from the exact solution of the CP obtained with conditional distributions *p*(*r_i_*|*D*) that are Gaussians (Dayan and Abbot, 2001; Carnevale et al., 2013) and not skew normals like for the threshold model (Eq.2). In fact, this CP approximation is generically valid when the activity-choice covariations are well captured by the linear dependence between the responses and the choice. Expanding Eq.7 in terms of *ρ_r_i__d* we get a polynomial approximation

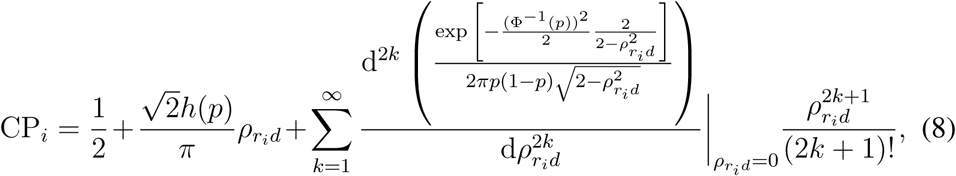

where *h(p)* is defined as

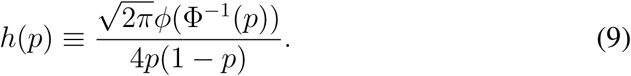

This factor is further discussed in the Results section. The expansion contains only odd order terms because of the symmetry of CP – 0.5 with respect to the sign of *ρ_r_i__d*. This explains why Haefner et al. (2013) found that the linear approximation was accurate for a wide range of *ρ_r_i__d*. Values, since the contribution of 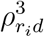. only starts to be relevant for intermediate to high correlations. Up to order 3 we have

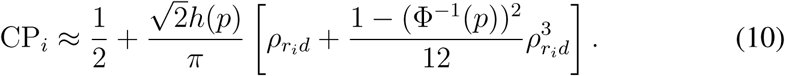

Since Φ^−1^(0.5) = 0, for the choice ratios for which |1 − (Φ^−1^(*p*))^2^| < 1 the third order term makes a smaller contribution than for the uninformative case. This is true for 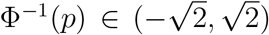 which leads to *p* ∈ (0.08,0.92). This means that the linear approximation is expected to be an even better approximation in this range than for *p* = 0.5. Furthermore, for (Φ^−1^(*p*))^2^ < 1 the third order contribution is positive, so that for the choice ratios fulfilling this constrain, *p* ∈ (0.16,0.84), the linear approximation is expected to underestimate the CP.

### 3.2 Derivation of the CTA for the threshold model

Following the work of Celebrini and Newsome (1994) as well as Britten et al. (1996), many studies have used the CP as a measure to characterize activity-choice covariations, and computational models have also focused on the CP (e. g. Cohen and Newsome, 2009; Haefner et al., 2013). A contribution of this work is to show that Choice Triggered Averages (CTAs) can provide complementary and often more useful information about the nature of activity-choice covariations. The mean response of cell i in trials where *D* = 1 is made is:

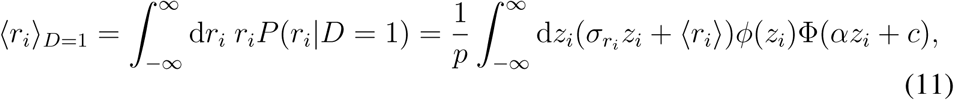

where the last equality holds for *P*(*r_i_*|*D* = 1) as in Eq.4. The CTA is defined as the difference of the mean responses for each choice, and using the integral equality of Eq.6 for *q* = 1,*n* = 1 into Eq.11, we have

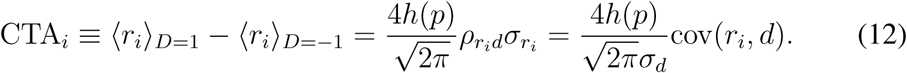

Furthermore, the fact that *D* is a binary variable, without any other assumption about the distribution of the responses, implies that

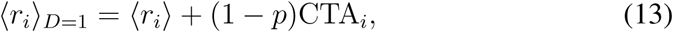

and similarly for 〈*r_i_*〉*D*=−1 substituting 1 − *p* by *−p.* That is, it is always the conditional mean of the less likely choice the one that contributes more to the CTA. It is straightforward to see the relation between the CTA in Eq.12 and the linear approximation of the CP (order one in Eq.10). Further details on their relation are discussed in the Results section.

### 3.3 Time-dependent analysis of CPs and CTAs

Above we have derived the CPs and CTAs without specifying for which interval the cell responses are estimated. In practice, a certain time window needs to be selected. In particular, for a time-resolved analysis responses are estimated in a sliding window to study the temporal changes of activity-choice covariations (e.g. Nienborg and Cumming, 2009). The strength of the influence of different sources of covariation is expected to change over the trial time. For example, serial dependencies across trials are expected to affect more the responses early in the trial, and on the other hand feedback signals containing categorical information about the choice can only exist after the decision. We now consider the time-resolved calculation of the measures. Consider a sliding window [*t_k_, t_k_* + *T*]. Following Eq.12, the CTA for *t_k_* is:

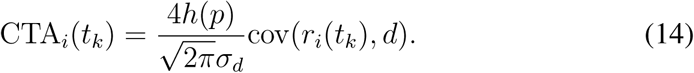

From the factors that determine the CTA only the covariance cov(*r_i_*(*t_k_*), *d*) is time-dependent. The choice ratio p, as well as the variance of the decision variable, are common to all *t_k_* because they are determined as a result of how all the sensory responses from different cells and time intervals are combined to generate d. Accordingly, if the responses result from the sum of different components each potentially associated with a different source of activity-choice covariation, then these contributions are linearly added in the CTA. For the CP, using the linear approximation from Eq.10, we have:

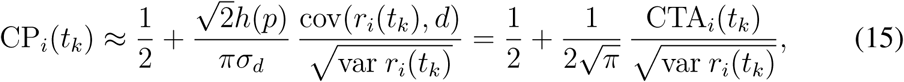

hence the CP temporal changes correspond to the relative changes of the CTA with respect to the changes in the response standard deviation.

### 3.4 Models of the different sources of activity-choice covariation

The derivations above provide an analytical formulation of the CP and CTA which does not make any assumptions about the sources that generate the activity-choice covariations. In this work we examine simple models representative of different sources and derive concrete expressions of the measures in each case. These models extend the feedforward linear model used by Haefner et al. (2013) by further considering feedback influences. They can approximate more general modulations of the responses by top-down signals (see Results for details) and are mathematically tractable, leading to close-form expressions that allow us to identify qualitative differences between how, for different sources of covariation, the measures depend on the model parameters.

We use Eq.12 to calculate the CTAs and the first order of Eq.10 to calculate the CPs. Accordingly, to characterize the measures all we need is to express in terms of the model parameters the following quantities: For the CTA, we need to derive the choice ratio p, the covariance cov(*r_i_*, *d*), and the variance *σ_d_.* For the CP, we also need the variance of the responses var *r_i_*. The choice ratio is calculated from Eq.3. For linear models, the variances and the covariance are obtained straightforwardly from their definitions. A detailed derivation can be found in the Supplementary material *S*1 of Haefner et al. (2013) for the feedforward model, and we proceed analogously for the new generic models incorporating feedback signals. Accordingly, in the Results section we directly present the final expressions for the CPs and CTAs obtained for each generic model. Furthermore, we also study some more concrete models, representative of a certain source of covariation. We now provide details on these concrete models.

#### 3.4.1 The neuron-antineuron feedforward model

The neuron-antineuron model is a subtype of feedforward threshold model in which it is assumed that the decision variable is constructed from the activity of two different pools of neurons, labeled as neurons and antineurons in relation to their response properties (Shadlen et al., 1996). Briefly, the neurons are more responsive for stimuli compatible with *D* = 1, and oppositely for the antineurons. The CP for this model was found to be mainly unsensitive to the distribution of the feedforward weights as long as they have opposite sign for the two pools. Furthermore, the CP is mainly unsensitive to the responses variance, and is determined by the difference between the average noise correlations between neurons within the same pool (*ρ_||_*) and noise correlations between neurons of different pools (*ρ_⊥_*). These results were analytically confirmed by Haefner et al. (2013). In particular, considering that all cells have equal variance and that each of the two types of correlations is homogeneous across cell pairs, in the limit of large pools the CP depends only on Δ*ρ* ρ *ρ*_||_ − *ρ_⊥_* (see equation S 1.14 of Haefner et al., 2013). However, for the time-dependent case (Eq.15), with these same assumptions and approximating the responses as independent in nonoverlapping time windows covering the trial time (for a fixed stimulus), we find:

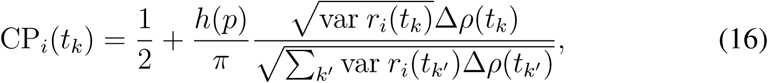

where *k*′ indexes the nonoverlapping windows. The denominator is the same for all times and corresponds to *σ_d_.* For a non time-resolved analysis with a single window, this expression of CP_*i*_(*t_k_*) reduces to the solution of Haefner et al. (2013), which only depends on Δ*ρ*. In the time-resolved case, temporal CP changes depend on the balance of the changes in Δ*ρ* and the variance, as we illustrate in the Results section.

#### 3.4.2 CTAs for a decision-related attention feedback model

Here we describe in detail a model of decision-related feedback through attention mechanisms (Maunsell and Treue, 2006). For a cell i with an unmodulated tuning function *f_0_*(*s − s_i_*), characterized by the preferred stimulus *s_i_,* attention leads to a modulated tuning function *f* (*s* − *s_i_*; *s_att_ − s_i_*) by introducing a multiplicative gain *g_i_*(*s_att_ − s_i_*), where *s_att_* is the attended stimulus. We consider this modulation to act as a decision-related feedback assuming that once the decision is made the attended stimulus is determined by the internal estimator of the presented stimulus, which is associated with *d*.

For illustration purpose, we more concretely conceive an orientation discrimination task in which the animal has to decide if the orientation of a bar has an angle higher or lower than a reference one. After the decision is made, the angle attended (*φ*_att_) is the predicted angle, estimated by *d*. Accordingly, for an unbiased internal estimator, the average predicted and attended angle coincides with the angle presented (*φ*). That is, 〈*φ_att_*〉 = φ. However, the trial-to-trial variability in *d* and hence in the predicted angle, leads to variability in *φ_att_* that influences the neural responses. This influence is the source of the activity-choice covariation. Linearly approximating this variability, the responses can be modeled as:

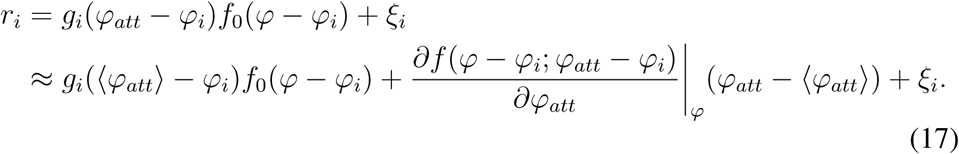

To implement the model, we follow Ecker et al. (2016) and use Von-Mises functions (Amirikian and Georgopulos, 2000) for the tuning curves. In particular, we take *f*_0_ (*φ* − *φ_i_*) = *q*exp[*κ* cos (*φ* − *φ_i_*)] and the gain as *g_i_*(*φ_att_*− *φ_i_*) *=* exp[*g* cos(*φ_att_* − *φ_i_*)]. Using these functions for the neural responses, and considering 〈φ^*att*^〉 = φ, leads to

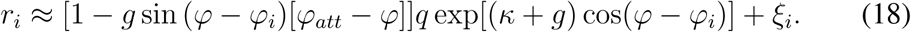

If *g* = 0, the responses are determined only by *f*_0_(*φ* − *φ_i_*). In general, the mean responses are

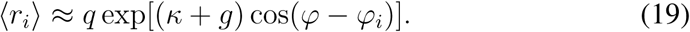

Given Eq.12, considering that the decision variable *d* provides an internal estimation of the angle, we can calculate the CTA in terms of cov(*r_i_*, *φ*_*att*_). In particular, from Eq.18

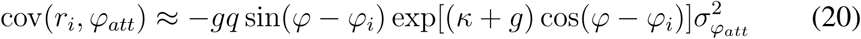

and

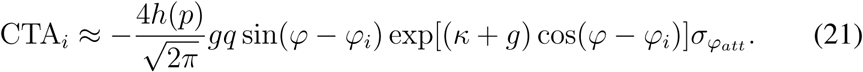

In the Results section we examine in detail the form of this CTA.

## 4 Results

We will present our results in several stages, progressing from the most general insights to specific applications. First, we present relationships for CPs and related measures of activity-choice covariation which hold independently of whether they are caused by feedforward (FF), feedback (FB), or a mixture of both kinds of signals. Second, we apply our formulas to plausible simple causal models of sensory responses and behavioral choices and derive the explicit dependence of CPs and CTAs on the parameters of interest in those models. In particular, we consider different sources of activity-choice covariation, comprising bias, internal states, and decision-related feedback signals. We then examine more specific scenarios for which our derivations are useful. First, we compare the time course of CPs and CTAs within a trial when the contributions to the covariation change over time. Second, we indicate the signatures in the CPs and CTAs of previously proposed computations that involve FB signals, like predictive coding and Bayesian inference. Finally, as a concrete example of discrimination between feedforward and feedback sources, we compare the CTAs derived for a model of decision-related attention feedback signals and for a feedforward model with optimal readouts.

### 4.1 General properties of activity-choice covariation measures

At the most basic level, any decision-related signal, independently of its origin, leads to an activity-choice covariation between the neural response (activity), *r*, and the psychophysical choice (decision), *D*. In the context of tasks involving two choices which we will consider here (comprising discrimination and detection tasks), this covariation is completely captured by the difference between the neural response distributions conditioned on the behavioral choice of the subject, *p*(*r*|*D* = −1) and *p*(*r*|*D* = 1). Among the possible measures to quantify the activity-choice covariation by comparing these two distributions, two related measures are choice probability (CP) and choice-triggered average (CTA):

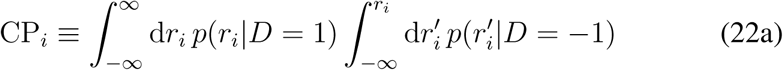

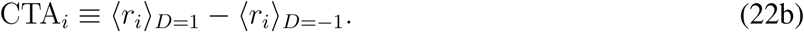

Intuitively, the CP is defined as the probability that a random sample from the distribution for choice 1 trials is larger than a random sample from the distribution for choice –1 trials (Britten et al., 1996; Parker and Newsome, 1998; Nienborg et al., 2012). It is 0.5 if both distributions are identical and goes to 0 or 1 as they are more and more separated. The CTA is defined as the difference between the mean response for trials of each choice (Haefner, 2015). It is zero when both distributions are equal. The CTA captures the linear dependencies between responses and choice and, given the binary nature of choice *D*, it is directly proportional to the covariance between them: CTA_*i*_ = 2cov(*r_i_*, *D*)*p*(*D = 1*)*p*(*D =* −1)1.

Furthermore, for small covariations (CPs close to 0:5) using the same linear approximation as in Haefner et al. (2013) one can show (see Methods) that:

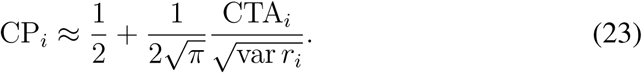

In this case the CP can be interpreted in relation to the response difference ‘normalized’ by the response variability (or the CTA of the *z*–score of the responses). Importantly, this relationship holds regardless of the causal sources of response-choice covariations, i.e. whether it is due to feedforward signals, feedback signals, or both, because it is agnostic regarding the origin of the dependence between *r_i_* and *D* (Fig. 1A). This relation between CPs and CTAs also shows that, when several independent sources contribute to the activity-choice covariation, the total CTA linearly adds the different covariance contributions from different sources, while linear additivity does not hold for the CP due to the denominator. For example, if *r_i_* = *r*_*i*_^(1)^ + r_*i*_^(2)^, then CTA_*i*_ = CTA_*i*_^(1)^ + CTA_*i*_^(2)^, where CTA_i_^(1)^ refers to the CTA that would be obtained if measuring only *r*_*i*_^(1)^, and analogously for CTA_*i*_^(2)^.

**Figure 1:**
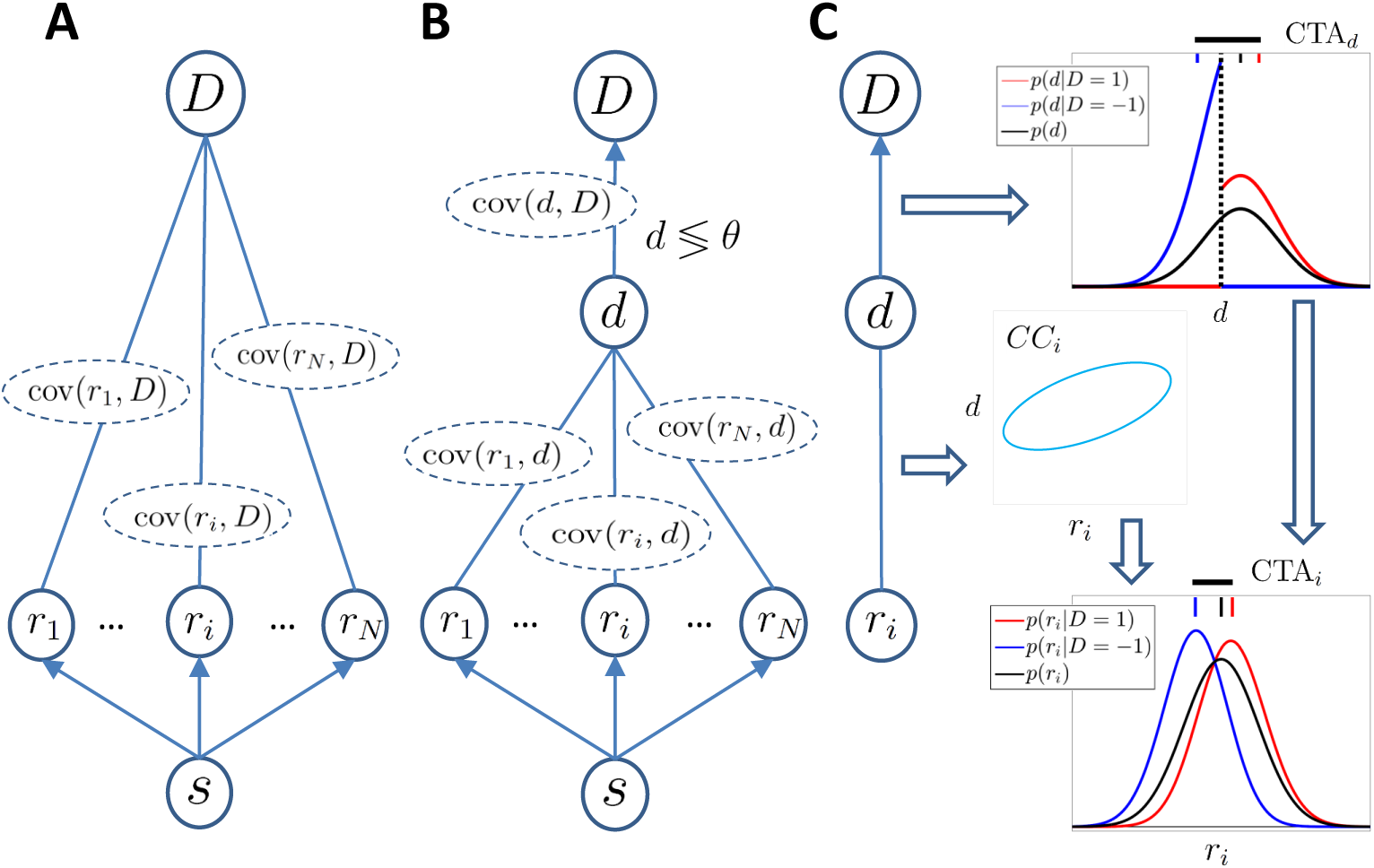
A general framework to analyze activity-choice covariations independently of the source of the decision-related signals. **A)** Representation of a completely agnostic covariations model: A population of sensory neurons encodes a stimulus s and activity covaries with the behavioural choice *D*. The arrows linking the stimulus and the responses indicate a causal influence. The undirected edges linking the responses and the choice indicate that we are agnostic about the source of the dependence reflected in the covariance cov(*r_i_*, *D*) between the activity of cell *i* and the choice *D*. **B**) Threshold model in which a continuous decision variable *d* mediates between the responses and the choice and represents an internal sensory stimulus estimator. The decision is made by comparing *d* with a threshold *θ*. **C**) Decomposition (characteristic of the threshold model) of the covariance between the response *r_i_* and the choice *D* in terms of the separated dependencies between the response and the decision variable *d*, and between *d* and the choice. The threshold mechanism (vertical dashed black line) dichotomizes the space of d, resulting in a difference between the mean of the conditional distributions associated with *D* = ±1 (red and blue vertical top bars, respectively). This difference is quantified by CTA_*d*_ (horizontal thick black line) and propagates to the Choice-Triggered Average (CTA_*i*_) when some correlation CC_*i*_ exists between *d* and *r_i_*.

So far we remained agnostic with respect to the mechanism relating sensory responses and behavioral choice. Now we adopt the common assumption that *D* is mediated by a continuous decision variable *d* that is an internal estimate of the sensory stimulus. We assume that *D* = 1 if the decision variable *d* exceeds an internal threshold *θ*, otherwise *D* = −1 (Gold and Shadlen, 2001, 2007). This allows us to separate the relationship between *r* and *D* in terms of the relationships between *r* and *d*, and *d* and *D* (Fig. 1B). In particular, the correlation coefficient between activity and choice is factorized such that corr(*r_i_*, *D*) = corr(*r_i_*, *d*)corr(*d*, *D*). Given the direct relation between CTA_*i*_ and the covariance cov(*r_i_*, *D*) we find

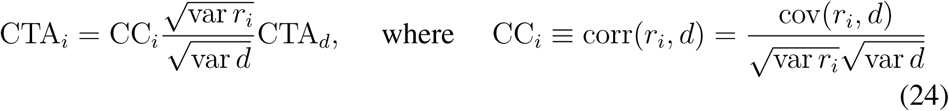

is the correlation coefficient between the sensory response and the decision variable termed Choice Correlation (Pitkow et al., 2015), and CTA_*d*_ is the difference of the means of the (unobserved) decision variable for the two choices. Eq.24 describes how activity-choice covariations are determined in the threshold model of Fig. 1B: The threshold mechanism dichotomizes the space of the decision variable into the two choices producing the CTA_*d*_, which propagates to CTA_*i*_ (Fig. 1C) given the specific correlation CC_*i*_ of the activity with the decision variable.

The threshold model is still agnostic with regard to the causal mechanisms introducing dependencies between *d* and the responses. We now add another assumption, namely that the decision variable is determined by the combination of the responses of a large population of neurons so that its distribution is well approximated by a Gaussian distribution. With this Gaussian assumption the dichotomization of the space of *d* is completely specified by the *choice ratio p,* i. e. the probability of choosing *D* = 1, which can be estimated from the ratio of the number of trials with each behavioural choice. Accordingly, the term 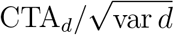 of Eq.24 can be expressed in terms of only *p*. In particular, we define a factor *h*(*p*) as the ratio of 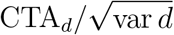 for a certain *p* with respect to its value for the uninformative stimulus (*p* = 0.5). Under gaussianity

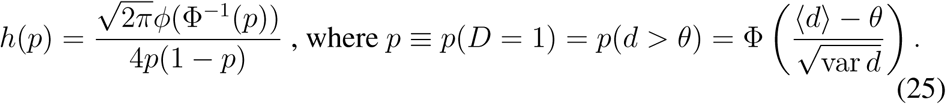

Here, *ϕ*(*x*) is the density function of a zero-mean, unit variance, Gaussian distribution, and Φ^−1^ is the corresponding inverse cumulative density function. 〈*d*〉 is the mean of the decision variable. By construction, *h*(*p*) = 1 for p = 0.5. Finally, plugging Eq.25 into Eqs. 23 and 24, the CTA and CP are expressed as

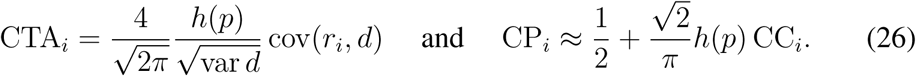

Note that the CTA_*i*_ and CP_*i*_ depend on the specific response properties of cell *i* only through cov(*r_i_*, *d*) and CC_*i*_, respectively. For the uninformative stimulus the CP of Eq.26 recovers the linear approximation derived in Haefner et al. (2013). The factor *h*(*p*) depends on the choice ratio and var *d* is also a factor common to all cells. Although generally the decision variable *d* cannot be directly observed, var *d* can be estimated based on the psychophysical threshold, fitting a psychometric function to the Gaussian expression of the choice ratio in Eq.25 (Pitkow et al., 2015).

Our theoretical approach starting from a totally agnostic model and sequentially incorporating more specific assumptions differs from previous theoretical studies characterizing CPs (e.g. Shadlen et al., 1996; Haefner et al., 2013) which assumed a feedforward model of activity-choice covariations (Fig. 2A). This approach provides three main new insights: First, the relationships in Eqs. 23-26 hold *regardless of the source* of the covariation - whether they are due to feedforward or feedback influences, common modulators, or some combination. This will allow us to study models with other causal architectures associated with non-feedforward sources of covariation (Fig. 2B). Fig. 2C summarizes how CP, CTA, and CC are interconnected. Second, the appreciation of CTAs. Even for a range of CP values for which the approximation that connects CPs and CTAs is not accurate, CTAs stand on their own as a characterization of activity-choice covariations, which capture only linear differences. As we will see, this reduced, more specific, sensitivity can in fact simplify the dependence on the parameters that identify a certain connectivity structure and help to discriminate between different potential sources of covariation. Third, the characterization of the dependence of CPs and CTAs on the choice ratio *p* by means of the multiplicative factor *h*(*p*). Considering this factor generalizes previous work that assumed the special case of *p* = 0.5, which is usually violated in practice (e.g. for trials with an informative stimuli).

**Figure 2:**
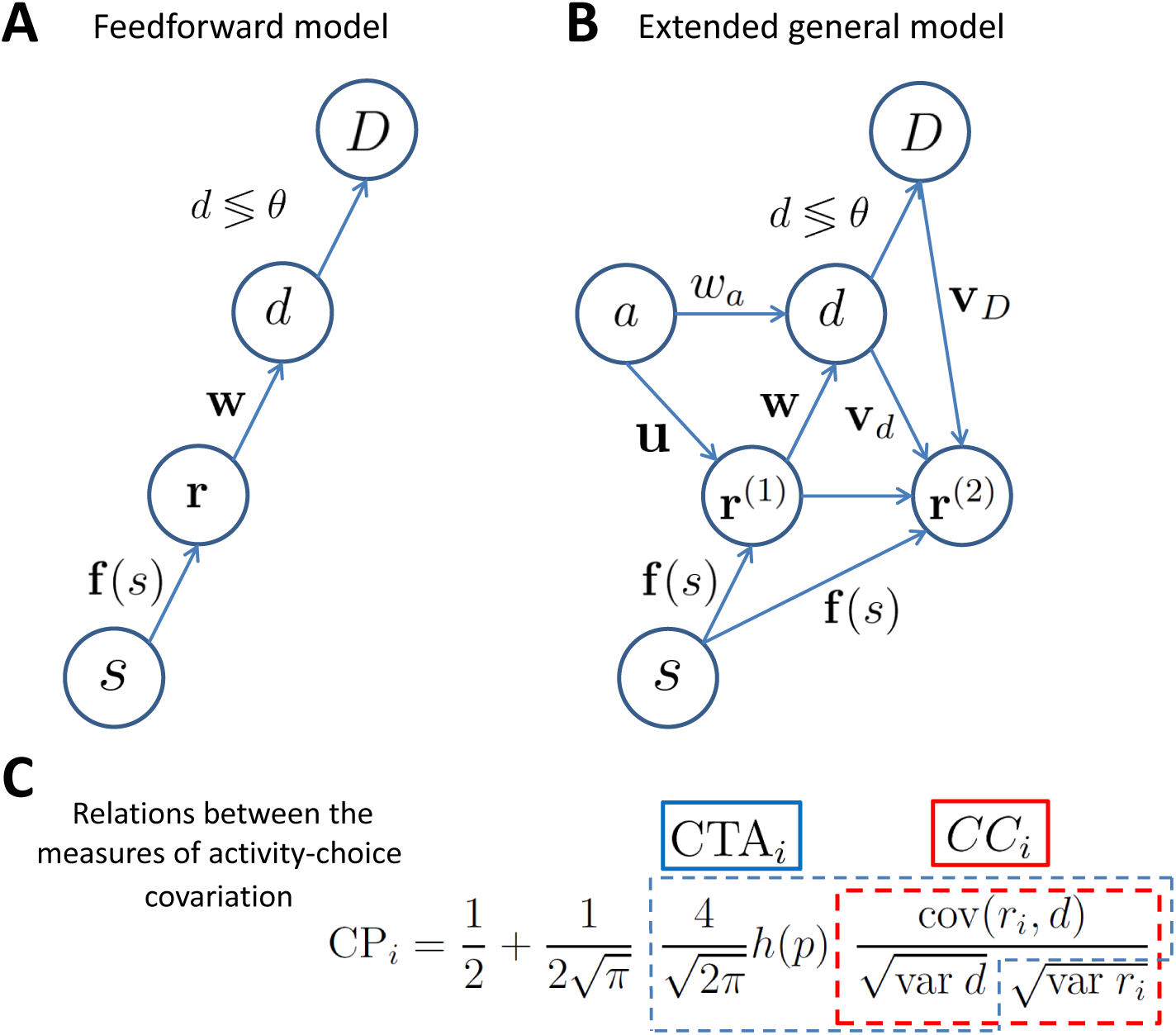
General models and measures to represent and quantify activity-choice covariations independently of their origin. **A**) Connectivity structure assumed for the traditional pure feedforward model. **B**) General architecture combining different sources of activity-choice covariation such as internal states (*a*) and decision-related feedback. In the main text we provide details of how the responses and the decision variable are generated for subcases focusing on different sources of covariation. r^(1)^ and r^(2)^ represent pre and post decision sensory responses. **C**) A graphical summary of the relation between the choice probability CP_*i*_, choice-triggered average CTA_*i*_, and choice correlation CC_*i*_. The complete picture of how these measures are interconnected is provided by Eqs. 23-26.

In the classical approach to examine activity-choice covariations, CPs are only estimated from ambiguous (i.e. zero-signal, or uninformative) trials (Britten et al., 1996). Attempts to more efficiently use all trials correct the influence of the stimulus to estimate a single CP value (Nienborg and Cumming, 2009; Kang and Maunsell, 2012). The generalized solutions introduced above for the case of informative stimuli indicate that changes in the stimulus informativeness may have an effect on the CP and CTA in two different qualitative ways. First, for the threshold model there is an intrinsic dependence on the stimulus through the multiplicative factor *h*(*p*) because the stimulus signal affects the choice ratio (Eq.25) either by the effect of its mean on 〈*d*〉 or by the effect of its trial-to-trial variability on 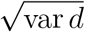. Second, the CP and CTA may further depend on the stimulus signal through the stimulus dependence of the choice correlation CC, for example if noise correlations are stimulus dependent. Separating these two types of effects of stimulus informativeness is necessary to more efficiently use data from all signal levels to study activity-choice covariations. One possibility, in line with previous approaches aiming to obtain a single measure from all trials, is to account for the factor *h*(*p*) to average the CP values obtained for different levels of stimulus signals. As an alternative, we here propose to study the CPs and CTAs as a function of the stimulus informativeness to gain new insights about the mechanisms originating the activity-choice covariations. As we will study below, different sources of covariation can be discriminated from the type of stimulus dependencies their produce between *r_i_* and d. These dependencies are reflected in the CP and CTA but we need to separate them from the intrinsic dependence on the stimulus signal that exists through *h*(*p*). Accordingly, we now proceed to characterize the shape of *h*(*p*) and consider how this factor may change when the gaussianity assumption does not hold. Furthermore, for the CP we compare its dependence on *h*(*p*) for the linear approximation (Eq.26) with an exact CP solution derived assuming joint gaussianity for *r_i_* and *d* (see Eq.7 in Methods for the form of this exact solution).

Fig. 3A compares, as a function of the choice correlation CC, an exact solution of CP_*i*_ (Eq.7) and the linear approximation shown above (Eq.26). For both the case of an uninformative stimulus (*p =* 0.5) and a highly informative stimulus (*p* = 0.9) the approximation is good for the range of CP values usually found experimentally (0.2 − 0.8). Fig. 3B shows CPs as a function of the choice ratio for three different values of the CC. To focus on the shape of *h*(*p*), we choose the CC to be invariant with the choice ratio (i.e. stimulus-independent) so that the CP reflects the dependence of *h*(*p*) on *p*. For *p* ≠ 0.5, *h*(*p*) > 1 and the CP increases symmetrically for *p* departing from 0.5. This can be understood intuitively, for example considering a sensory neuron feedforwardly contributing to the decision variable: A highly informative stimulus induces signal-dominated neural responses, so that *d* most likely lies on the side of the threshold compatible with the sensory stimulus presented (e.g. *D* = 1) and leads to *p* close to 1. This means that *p*(*r_i_*|*D* = 1) is similar to *p*(*r_i_*), since most responses correspond to *D* =1, and hence its conditional mean is also close to the unconditional one. On the other hand, the opposite choice is made only for trials with a substantial and contradictory departure of the neural responses from the signal-driven mean response. Accordingly, the distribution *p*(*r_i_*|*D* = −1) contains responses that lie in the tail of *p*(*r_i_*). As p goes to 1, the mean of *p*(*r_i_*|*D* = 1) converges to the unconditional mean, while the departure of the mean of *p*(*r_i_*|*D* = −1) from the unconditional mean becomes increasingly large, resulting in an increasing difference between the two (Fig. 1C, and see Eq.13 in Methods).

**Figure 3:**
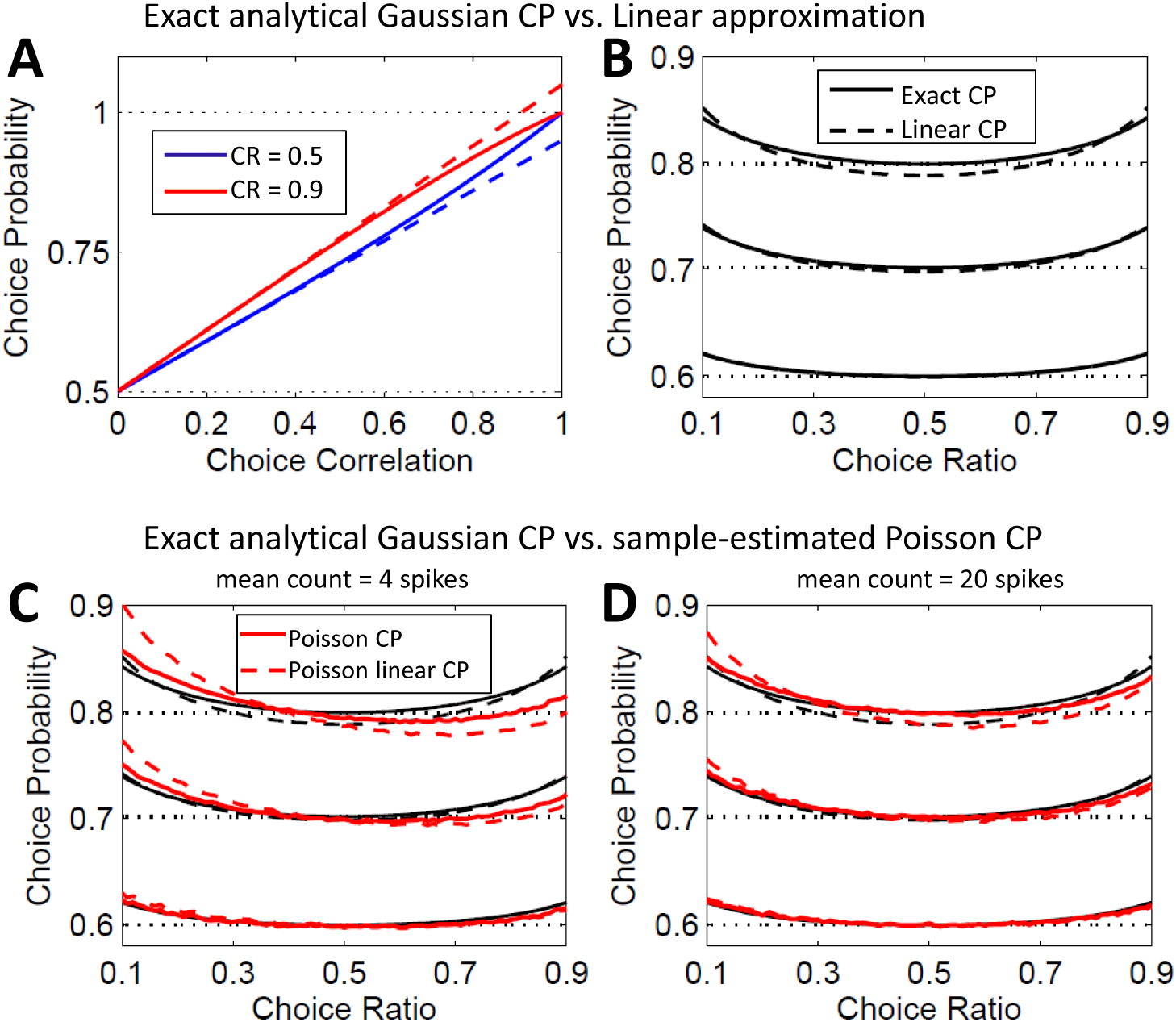
Choice probabilities calculated from the threshold model in the presence of informative stimuli. **A)** Comparison, as a function of the choice correlation, of the exact solution of the CP (solid lines, Eq.7) and its linear approximation (dashed lines, Eq.26). Results are shown for two values of the choice ratio p, 0.5 and 0.9. **B)** Same comparison but with the CP as a function of the choice ratio. Results are shown for three values (horizontal dotted lines) of the exact CP for the uninformative case p = 0.5, each determined by a different CC value. Again solid lines represent the exact solution and dashed lines the linear approximation. **C)** Comparison of the exact Gaussian CP solution with the estimations obtained from simulated Poisson responses with low spike counts. Black lines reproduce the analytical results already shown in panel B. Red lines correspond to the estimated CP from simulated Poisson data. Estimation is based directly on the CP definition (red solid line, Eq.22), to be compared with the exact Gaussian solution (black solid line), and on the CP linear approximation (red dashed line, Eq.23), to be compared with the linear Gaussian approximation (black dashed line). **D)** Same as C) for Poisson responses with higher counts.

The shape of *h*(*p*) indicates that an increase of CPs with the choice ratio provides evidence compatible with the decision making threshold mechanism. However, the influence of the stimulus signal through *h*(*p*) is small for a substantial range of the choice ratio around *p* = 0.5 and its characterization may require large numbers of trials (i.e., chronic recordings or pooling across many neurons). On the other hand, this relatively flat dependence justifies the combination of trials with different signal strengths to calculate a single CP value per neuron (so-called ‘grand choice probabilities’, Nienborg et al., 2012), if no further stimulus dependence exists through the CC.

Fig. 3C-D compare CPs and CTAs obtained analytically with those estimated from simulations of a feedforward model. The decision variable is obtained as the sum of a Poisson component, modeling the contribution of cell i and other cells correlated to it, and of a gaussian component, modeling the contribution of a large number of neurons uncorrelated to cell *i*. The relative weight of these two contributions determines the CC. CTAs have been transformed to CPs with Eq.23 so that also the accuracy of this linear approximation can be checked. For these transformed CTAs (dashed lines), the comparison of the analytical (black) and estimated values (red) exclusively reflects the accuracy of *h*(*p*) derived under gaussianity. For CPs, the comparison of analytical (solid black) and estimated values (solid red) reflects the accuracy of the whole exact CP solution derived under gaussianity.

The factor *h*(*p*) is well approximated by its gaussian form except when *d* results from the combination of activity of a small population, or of a population of highly correlated neurons, in a way that the central limit theorem that justifies the Gaussian approximation is less accurate. In our simulations these departures from the analytical values occur for high CCs, especially when the firing counts are low. This can be understood from how we generated the data: to obtain high CCs for cell *i* we make the weight of the Poisson contribution to *d* higher. For high counts, the Poisson contribution is itself well approximated by a Gaussian, but for low counts it is not and thus the distribution of *d* departs from a Gaussian. In general the spike counts of a cell do not indicate how good the gaussian approximation of *h*(*p*) is, since this is a factor common to all cells. The form of the departures from the analytical values can also be qualitatively understood: symmetry with respect to *p* = 0.5 is broken because of the effect that for Poisson responses the constraint to zero or positive counts has on how *p*(*r_i_*|*D =* 1) and *p*(*r_i_|D =* −1) can mutually differ. For *p* > 0.5 the mean of *p*(*r_i_|D* = −1) should be shifted to smaller counts and predominantly contribute to the CTA, since the conditional mean of the responses for the less likely choice differs more from the unconditional mean (Fig. 1C). However, this shift is constrained by counts being zero or positive, especially for low rates. Overall, Fig. 3C-D show that the Gaussian approximation for *d* is robust and Eq.25 can be used to approximate *h*(*p*) and to correct for the influence of the choice ratio on CPs and CTAs. Only if the decision variable relied on a small number of neurons or on a larger but highly correlated population the approximation would be less accurate.

### 4.2 Activity-choice covariation on specific causal models

The derivations above provide an analytical formulation of the CP and CTA measures making as few assumption as possible about the sources that generate the response-choice covariation, in particular about the connectivity architecture linking the neural responses to the decision variable. In this section we will examine simple plausible models representative of different sources of response-choice covariations (Fig. 2B). This will allow for a better understanding of how different signals influence CPs and CTAs. First we present the results for models that consider feedforward and feedback influences separately and then we describe the combination of several influences into CTAs and CPs. For simplicity of presentation, we will explicitly indicate stimulus dependencies (e.g. CP(s)) only when needed. We will return to them in sections 4.4 and 4.5 because these stimulus dependencies can help to identify the sources of covariation.

#### 4.2.1 Pure feedforward model

First, we consider the classic FF encoding/decoding model (Fig. 2A, Shadlen et al., 1996; Haefner et al., 2013) in which a population of sensory responses, **r** = (*r*_1_,…, *r_n_*), is being read out by a decision area representing a decision variable 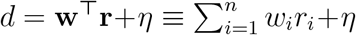, where w are the feedforward read-out weights and *η* represents some decision noise additional to the variability in the sensory responses (called ‘pooling noise’ in Shadlen et al. (1996)). The categorical choice *D* is made by comparing *d* to the threshold *θ.* We model the responses as *r_i_ = f_i_*(*s*) + ξ_*i*_, with means given by the tuning functions f(*s*) *=* (*f*_1_(*s*)*,…, f_n_*(*s*)) and covariance structure C_ξ_ of the neuron’s intrinsic variability ξ_*i*_. Note that **r** may comprise the responses of different cells but also the responses at different times, so that the covariance matrix C_ξ_ generically captures both the responses cross-correlations and autocorrelations. In general, the total measurable covariance matrix C may have other contributions apart from the intrinsic variability of the responses quantified by C_ξ_, for example due to trial-to-trial variability in the stimulus signal (see Supplementary Material *S*1 of Haefner et al., 2013, where the CP solution of the feedforward model is derived considering this stimulus variability). For simplicity, we do not include stimulus variability in our models and hence for this feedforward model we have that C = C_ξ_. Nonetheless, as we will see, internal variability, for example decision-related, may further contribute to C. From Eq.26 in this feedforward model

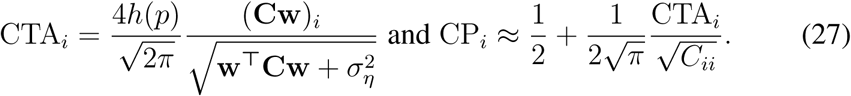

Note that *C_ii_ =* var *r_i_*. For *p =* 0.5, i.e. *h*(*p*) *=* 1, we recover the solution previously found by Haefner et al. (2013). In this feedforward model activity-choice covariations are explained as a population effect because the covariance (Cw)_*i*_ does not arise from the single influence on *d* of *r_i_* through its own weight *w_i_* (Shadlen et al., 1996; Haefner et al., 2013). This direct single influence is expected to be small when a large neural population determines the decision, but the covariance captures the contributions from all indirect connections due to crosscorrelations with the rest of the population. Given the form of the covariance (Cw)_*i*_, the estimation of feedforward weights requires estimating the covariance matrix C (Haefner et al., 2013). Accounting for *h*(*p*) allows one to pool the CPs from all signal strengths in order to infer the read-out weights w, instead of using only the CP calculated from zero-signal trials.

#### 4.2.2 A generic view of models with neural responses modulated by a decision-related internal variable

A characteristic of the feedforward model is that the responses are modeled as *r_i_* = *f_i_*(*s*) *+ ξ_i_,* with a component which is stimulus driven and a component considered as noise, basically containing any other source of variability of the responses. More realistically, the trial-to-trial variability of the responses is expected to be at least partly explained by changes in internal states (e.g. Masquelier, 2013; Ecker et al., 2014; Rabinowitz et al., 2015). These internal variables may have top-down effects on the neurons responses under study and also introduce variability in the choice through other pathways, hence producing some activity-choice covariation. Internal states also comprise the decision and choice variables themselves, which can modulate the responses through decision-related feedback.

To be more concrete, consider a neuron *i* whose mean response can be modeled as *f_i_*(*s*; *s*′) = *f_i−FF_*(*s*) + *f_i_*_−*FB*_(*s*; *s*′). Here *s* is the presented stimulus, and s' is an internal state, which usually cannot be controlled by the experimenter. The tuning curve has two components: *f_i_*_−*FF*_(*s*) models the feedforward response in the absence of any feedback signals and *f_i_*_−*FB*_(*s*; *s*′) represents the contribution of the feedback signal. The feedback can be multiplicative or additive depending on *f_i_*_−*FB*_ being or not proportional to *f_i_*_−*FF*_, and it is decision-related if *s*′ is correlated with the actual choice. To study the feedback effects in a tractable way we use a linear approximation considering that *s*′ fluctuates around its mean 〈*s′*〉. Accordingly, the responses *r_i_* are approximated as:

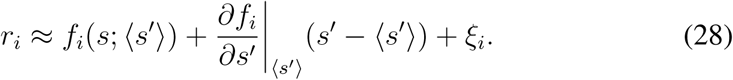

This constitutes the general form of a responses model which is linear in the contribution of the variability related to the internal variable *s*′. The derivative of the tuning curve with respect to *s*′ corresponds to a linear feedback weight. We will use the notation *f′_s_*_′_ to refer to this derivative. Since it is the variability of *s*′ what produces the activity-choice covariation induced by the internal variable, the CTA will be related to this derivative. The feedback also alters the structure of the covariance matrix of the responses by adding noise correlations caused by s^'^ as a common source of variability. In particular, 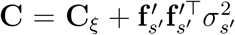, and thus the influence of feedback naturally introduces a relation between the structure of CTAs, noise correlations, and tuning properties. As we mentioned above, C generically includes zero and non-zero lagged cross-correlations as well as autocorrelations, and hence noise correlations shaped by the decision-related feedback can determine the structure of all these correlations (Wimmer et al., 2015; Haefner et al., 2016). Below we examine specific cases of this general model. To distinguish the feedback weights in the case of feedback from the decision variables themselves or any other internal variable, we will refer to these weights as *v_i_* and *u_i_,* respectively. In both cases they can be related to *f′_s′_* in agreement with Eq.28.

#### 4.2.3 Feedforward model including top-down signals from an internal variable

We first consider the same FF model while allowing a common input from an internal variable *a* to influence both neural responses **r** and the decision variable *d* (Fig. 4A). The internal variable can represent different mechanisms such as an expectation bias, cognitive memory effects that link choices across subsequent trials (e. g. Frund et al., 2014; Conen and Padoa-Schioppa, 2015), slow modulations in the level of excitability (e. g. Goris et al., 2014), or other sources of long memory autocorrelations in the neural responses that also introduce dependencies across trials (e. g. Nienborg and Macke, 2014). We model the influence of this common input on **r** and *d* by linear weights u and *w_a_*, respectively. Its direct influence on the decision variable *d* is captured by *w_a_,* that is, *d* = w^T^r + *w_a_a* + *η*. The response of neuron *i* is given by *r_i_* = *f_i_*(*s*) + *u_i_*(*s*)*a* + ξ*_i_*. Without loss of generality, we consider *a* with mean 〈*a*〉 and unit variance. As discussed above, u can depend on s and introduce both additive, multiplicative and intermediate influences of a on the responses. In comparison with the general model of Eq.28, here *a* corresponds to the state *s*′. The component of the response depending on has 〈*s′*〉 been absorbed in *f_i_*(*s*) to simplify the notation. We do not explicitly model the form of *f_i−FB_*(*s*; *s*′), and we generically assume that it determines the weights u through the derivative *f′_a_*. We obtain:

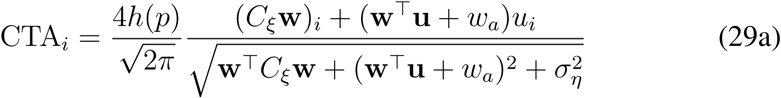

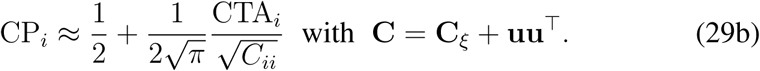

**Figure 4:**
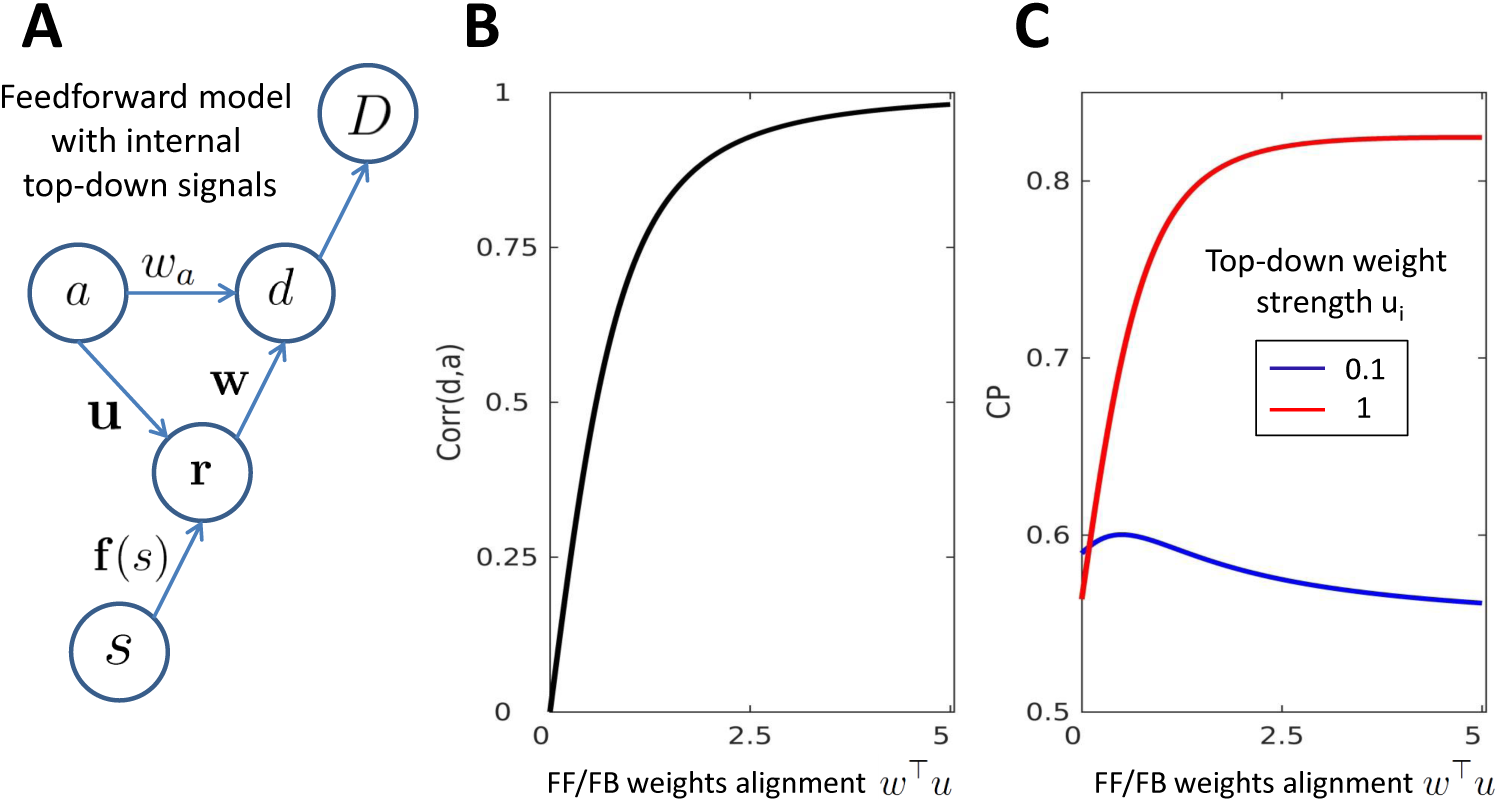
The effect of top-down influences from internal variables on activity choice covariations. **A**) Causal architecture of a model combining feedforward influences with top-down influences from an internal variable *a.* **B**) Correlation Corr(*d*, *a*) between the decision variable and the internal variable *a* as a function of the degree of alignment of the top-down and feedforward weights (w^T^u). This alignment determines the feedforward contribution of the top-down signal to the decision variable. **C**) Choice probability for this model (Eq.29) as a function of w^T^u for two different strengths of the top-down weight *u_i_* of cell *i*.

Changes in the mean top-down signal may introduce a choice bias and affect the measures only through *h*(*p*), either by altering the mean of the decision variable or the decision threshold *θ*. To understand how the variability in a influences the activity-choice covariations we consider first the subcases in which it affects only either the decision variable (**u** = 0) or the responses (*w_a_* = 0). In the first case, *a* only increases the decision variability that is unrelated to the responses **r** and hence decreases the covariations. In the second case, the internal variable increases the decision variability related to **r** by adding a new component to the responses. The new term uu^T^ in the measurable covariance matrix C captures the covariability added by the common input *a.* The top-down signal to the responses propagates in a feedforward way to the decision variable depending on the alignment between the feedforward and feedback weights, w^T^u. The CTA can be rearranged to recover the form of the purely feedforward model as in Eq.27. In particular, (*C*w)_*i*_ = (*C*_ξ_w)_*i*_ + w^T^u*u_i_* and w^T^*C*w = w*^T^C_ξ_*w + (w^T^u)^2^. Finally, when both u and *w_a_* are nonzero, these two influences coexist with a third that arises specifically from the effect of the internal variable as a common driver of the decision variable and the responses, as reflected in the interaction terms *u_i_w_a_* and w*^T^uw_a_* in the numerator and denominator of Eq.29 a, respectively.

Since the internal variable influences var *r_i_*, var *d*, and cov(*r_i_*, *d*), its net effect depends on its relative contribution to them. In particular, the degree of alignment of w^T^u determines how well the top-down signal feedforwardly propagates to the decision variable (Fig. 4B). If these weights are aligned the variability from the internal variable is transferred to d, increasing its variance var *d*. The effect on activity-choice covariations with a specific cell *i* depends on how sensitive the cell is to the internal variable variability. If *u_i_* is small, its responses are weakly driven by the internal variable and hence the increased variability in the decision variable is perceived mostly as noise, leading to a decrease in the CP (Fig. 4C). Oppositely, if *u_i_* is high, when the alignment increases the top-down signal simultaneously drives the cell responses and is propagated to *d*, and hence the CP increases. In Fig. 4B-C, Corr(*d, a*) and CP are calculated fixing all the model parameters and varying only w^T^u and *u_i_*. We change w^T^u while keeping constant var *r_i_* and thus the CTA profile of changes as a function w^T^u is the same as the CP profile shown in the figure.

#### 4.2.4 Effects of a post-decision feedback signal

While an internal variable commonly influences the responses and the decision variable, we now consider feedback signals from the decision variable itself that affect the responses *after the decision has been made.* Feedback signals have been invoked to explain activity-choice covariations in late trial time intervals (Nienborg and Cumming, 2009). The time at which the decision is made naturally splits the trial into two phases (Kiani et al., 2008). We denote the sensory response during these parts by r^(1)^ and r^(2)^, respectively. The analysis of activity-choice covariations for r^(1)^ corresponds to cases examined above. For the feedback phase we will consider two plausible cases separately. First, we assume that the feedback strength is proportional to the continuous decision variable *d* and thereby related to the degree of certainty in the decision (Fig. 5A). Second, we assume the feedback to originate from the categorical choice variable *D* and only depend on the choice itself (Fig. 5B). We talk about a feedback influence because we conceive **r** as the response of a neuron in a sensory area receiving a decision-related influence that modulates its sensory response. Alternatively, we can also consider decision-related influences for neurons located in decision-making areas or in downstream areas of the decision-making process, such that the decision-signal can account for a large part of the neuron’s trial-to-trial variance (e. g.Katz et al., 2016). While these cases are not represented by the causal architectures from Fig. 5A-B the derivations below also comprise them.

**Figure 5:**
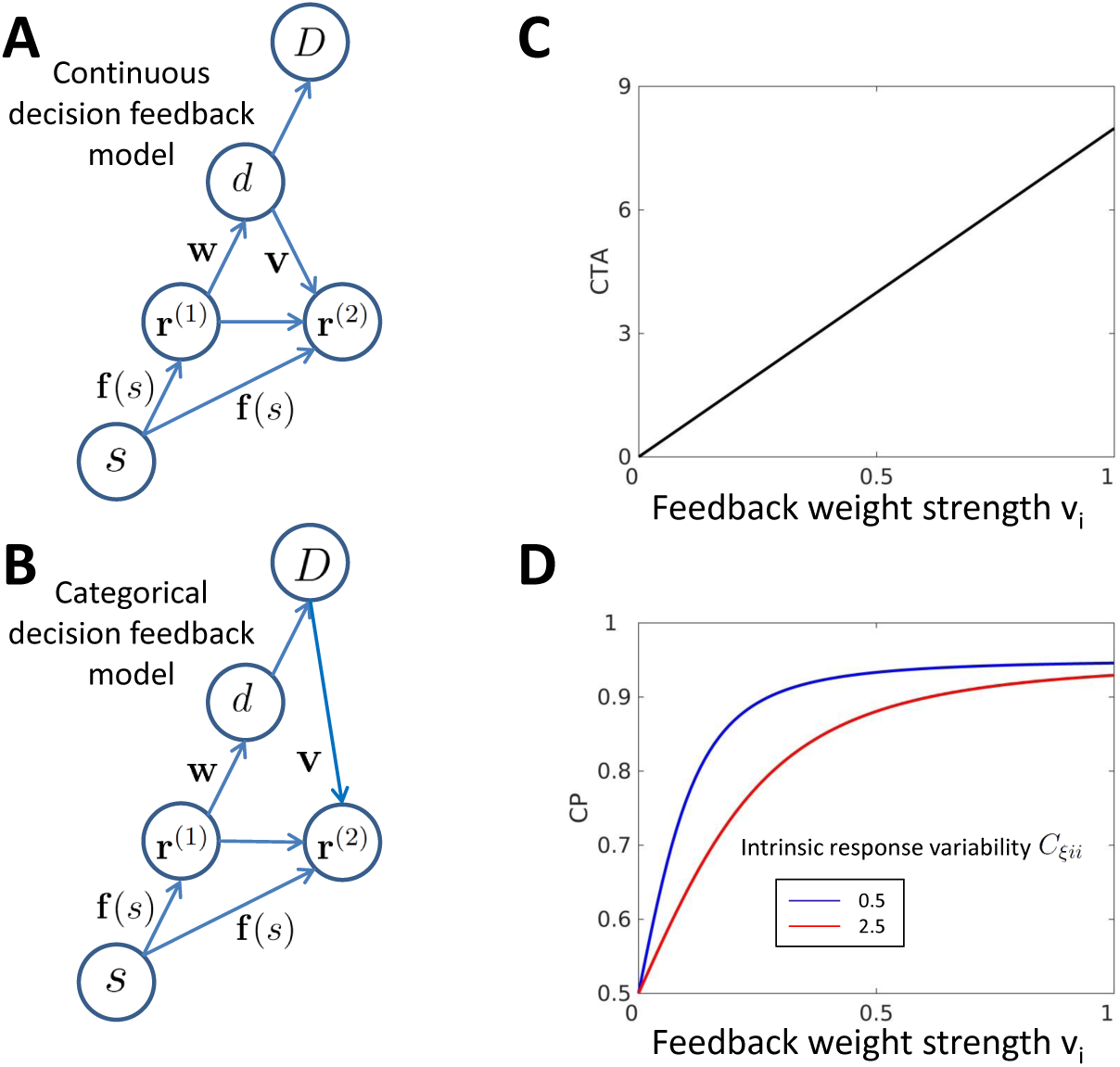
The effect of decision-related feedback on activity-choice covariations. **A**) Causal architecture of a model combining feedforward influences with postdecision feedback from the continuous decision variable. **B**) Causal architecture of a model combining feedforward influences with post-decision feedback from the categorical choice variable. **C**) Choice triggered average as a function of cell i feedback weight strength. **D**) Choice probability as a function of cell i feedback weight strength for two levels of intrinsic neural response variability, determined by the variance *C_ξii_*. CTAs and CPs are calculated for the pure continuous feedback model according to Eq.30.

##### Continuous feedback

We model the response of neuron *i* after the decision as *r_i_*^(2)^ = *f_i_*(*s*) + *v_i_*(*s*)*d* + ξ_*i*_, where v = (*v*_1_,…, *v*_n_) quantifies the strength of the influence of *d* on the sensory responses. The post-decision responses r^(2)^ do not affect the decision, in contrast to the previous responses r^(1)^. Accordingly, we also differentiate the covariance matrices C^(1)^ and C^(2)^. As for the internal variable *a,* we allow v to depend on the stimulus to model additive, multiplicative, and intermediate feedback. Furthermore, the model is general enough to consider that the feedback affects a different group of cells than those involved in the decision process. This is the case for cell *i* if *w_i_* = 0 and *v_i_* is nonzero. We obtain for the measures of activity-choice covariation:

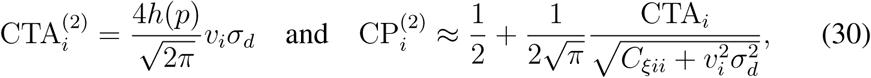

where the decision variable variance is 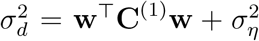 and the covariance matrix is 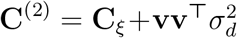, in contrast to C^(1)^ = C_ξ_. Accordingly, analogously to the internal variable a, decision-related feedback contributes to the measurable covariance with the term 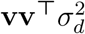. The asymmetry between the feedforward and feedback pathways is reflected in the form of the covariance *cov*(*r_i_,d*), which determines the CTA. In the feedforward case, there is a many-to-one relation between the responses **r** contributing to the decision variable and *d* and hence, as discussed above, it is well understood that covariations do not arise from the direct influence of a single cell on the decision, which is expected to be small, but from all indirect dependencies due to correlated activity with the rest of the population. In the feedback case, *d* can exert its influence directly on each neuron to produce activity-choice covariations. This one-to-one influence is reflected in the proportionality between the CTA_*i*_ and the feedback weight *v_i_* of cell *i* (Fig. 5C). This linear relation with the feedback weight is specific to the CTA. For the CP, the normalization introduces a nonlinear dependence on *v_i_*. In Fig. 5D we compare how CP increases as a function of the feedback weight for two different values of the intrinsic response variance. The CP rate of increase when *v_i_* increases depends on the response variance, and hence it is initially smaller when the intrinsic variance is high. Accordingly, how CP increases with feedback depends on its relative effect on the response variance and on cov(*r_i,_ d*).

##### Categorical feedback

In this scenario, we assume that the feedback signal only depends on the choice itself. Feedback associated with the categorical choice naturally leads to a bimodal distribution of the responses, since it introduces a discrete shift from one choice to the other. Accordingly, we model the responses as *r_i_* = *f_i_*(*s*)*+ v_i_*(*s*)*v_D_* + ξ_*i*_, where the feedback signal *v_D_* is separately distributed for *D = ±*1 as a Gaussian distribution with mean ±Δ_*D*_/2 and variance 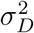. For Δ_*D*_ = 0, this signal does not depend on the choice and only adds unrelated noise to the responses. We find:

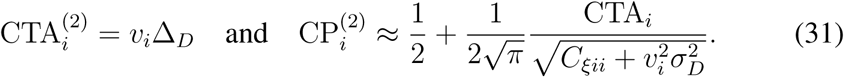

Like for the continuous feedback case, CTA_*i*_, but not CP_*i*_, is linearly proportional to the feedback weight *v_i_*. The qualitative difference between the categorical and continuous feedback is reflected in the dependence of the CTA on Δ_*D*_ instead of *σ_d_.* Accordingly, in contrast to the continuous feedback, the categorical feedback is unsensitive to the variability in the decision variable associated with the internal estimation of the stimulus. This difference can be exploited to discriminate between the two types of feedback, in particular by examining the influence that changes on *σ_d_* have on the CTA, for example manipulating the stimulus trial-to-trial variability to control the variability in the decision variable.

##### Combination of feedforward and feedback signals

So far we have considered the pure feedback models for r^(2)^ in isolation, as if any correlation between r^(2)^ and r^(1)^ was mediated by the feedback. However, the form of cov(*r_i_, d*) in the feedforward model, (Cw)_*i*_, indicates that a feedforward contribution is expected also for the second phase if the intrinsic variability of the responses during the two phases is autocorrelated. This is because separating the responses into r^(1)^ and r^(2)^ is equivalent to dividing the global covariance matrix C into the block diagonal parts C^(1)^, C^(2)^, and the (symmetric) block off-diagonal parts, C^(21)^. The feedforward term (C^(21)^w)_*i*_ can contribute to the covariation of 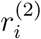 with the choice even if cells r^(2)^ do not determine the choice.

More generally, a mixture of feedforward and feedback contributions to the activity-choice covariations is expected when responses are estimated for a time interval that includes parts from both phases, before and after the decision time. If the decision time was fixed across all trials, the total CTA would be the sum of the respective CTAs of both phases, as described earlier. However, this time also varies from trial to trial and its variability may itself contribute to activity-choice covariations if, as expected, the decision time is associated with the amount of evidence accumulated. The CTA then has two contributions, one directly related to the decision time covariation with the decision and another determined by the average covariation of the responses with the decision for a fixed decision time. In this work we do not further consider the influence of decision time variability and we assume that either its contribution is small, so that the total CTA is approximately the sum of the feedforward and feedback CTA components, or covariations are quantified for time intervals in which either feedforward signals dominate (e.g. early in the trial) or feedback signals dominate (late in the trial) so that the equations of the pure feedforward or feedback models can be used, respectively.

#### 4.3 Complementary information of CTAs and CPs about temporal changes of activity-choice covariation

Above we have derived how the empirically observable quantities CTA, CP, and covariance C depend on the structure and parameters of the underlying models that lead to activity-choice covariations. Our derivations indicate the utility of the CTA as a measure related but complementary to the CP. The CP linear approximation in terms of the CTA and of the responses variance (Eq.23) establishes a link between these three measurable quantities, hence advocating for their joint analysis. In principle, given that the CP is determined by the two others, it would suffice to study the CTA and the variance to characterize activity-choice covariations. However, since the relation between these variables only strictly holds in the linear approximation, the CP may reflect other aspects of the covariation. Furthermore, calculating the CP is relevant if only to link the results to previous studies predominantly using this measure.

We illustrate the utility of this joint analysis in a particular case, namely the estimation of time-resolved CPs over the trial time. This temporal analysis, in combination with psychophysical kernels, has been used to evaluate the contribution of feedforward vs. feedback sources to activity-choice covariation (Nienborg and Cumming, 2009). We here show how the CTA and the response variance can provide complementary information in this type of analysis. The link between CPs and CTAs in the linear approximation indicates that CP changes across time correspond to CTA changes normalized by the variance of the responses (Eq.23, and see Eq.15 for details). Since this variance can also be time-dependent, the CPs can change even when the CTAs remain constant. For example, for a Poisson-like response the variability is proportional to the average rate, and the rate profile may change over the trial e.g. decaying after a transient increase locked to the stimulus onset. Indeed, Eq.23 shows that any combination of increased/decreased CP/CTA is possible, depending on the response variance.

Fig. 6A-B display some plausible temporal profiles of CP_*i*_ and CTA_*i*_ due to changes over time in the top-down weight *u_i_* (*t*), associated with the influence of an internal variable (blue), and changes in the feedback weight *v_i_*(*t*), for a continuous decision-related feedback (red). Since the CTA is not normalized and we are interested in its changes we here show the ratio of the CTA relative to its initial value at *t* = 0 (rCTA). Given Eq.26, this ratio directly corresponds to a ratio of covariances, since all the other factors that determine the CTA are constant over time. We are mainly interested in the trend of increase or decrease of the measures over time, while the exact shape of their temporal profiles is determined by the profiles of temporal change of the weights. Here we have modeled the changes in the top-down weight *u_i_*(*t*) as an exponential decay (∝ *e*^−*t*/*τ*^) and oppositely the changes in *v_i_*(*t*) as an exponentially-saturating increase (∝ 1 − *e*^−*t*/*τ*^), where *τ* determines the changes time-scale. These variations mimic the following plausible changes: For example, for serial dependencies across trials, the internal variable is expected to determine the neural responses mainly in the early part of the trial. Oppositely, decision-related feedback is expected to affect more a later phase of the trial. For the internal variable, we consider the measures derived in Eq.29 in their time-resolved form. If the internal variable varies slowly, in a longer time scale than the trial time, the measures have an analogous form, but with a time varying weight *u_i_*(*t*) in the numerator of the CTA. For the decision-related feedback, we use the continuous feedback model (Eq.30) and we further add feedforward activity-choice covariations to model the transition from a feedforward dominated trial interval to a feedback dominated interval that occurs when the sliding window is shifted towards the late trial time. We keep the feedforward contribution constant and change the feedforward/feedback relative strength by changing *v_i_*(*t*).

**Figure 6:**
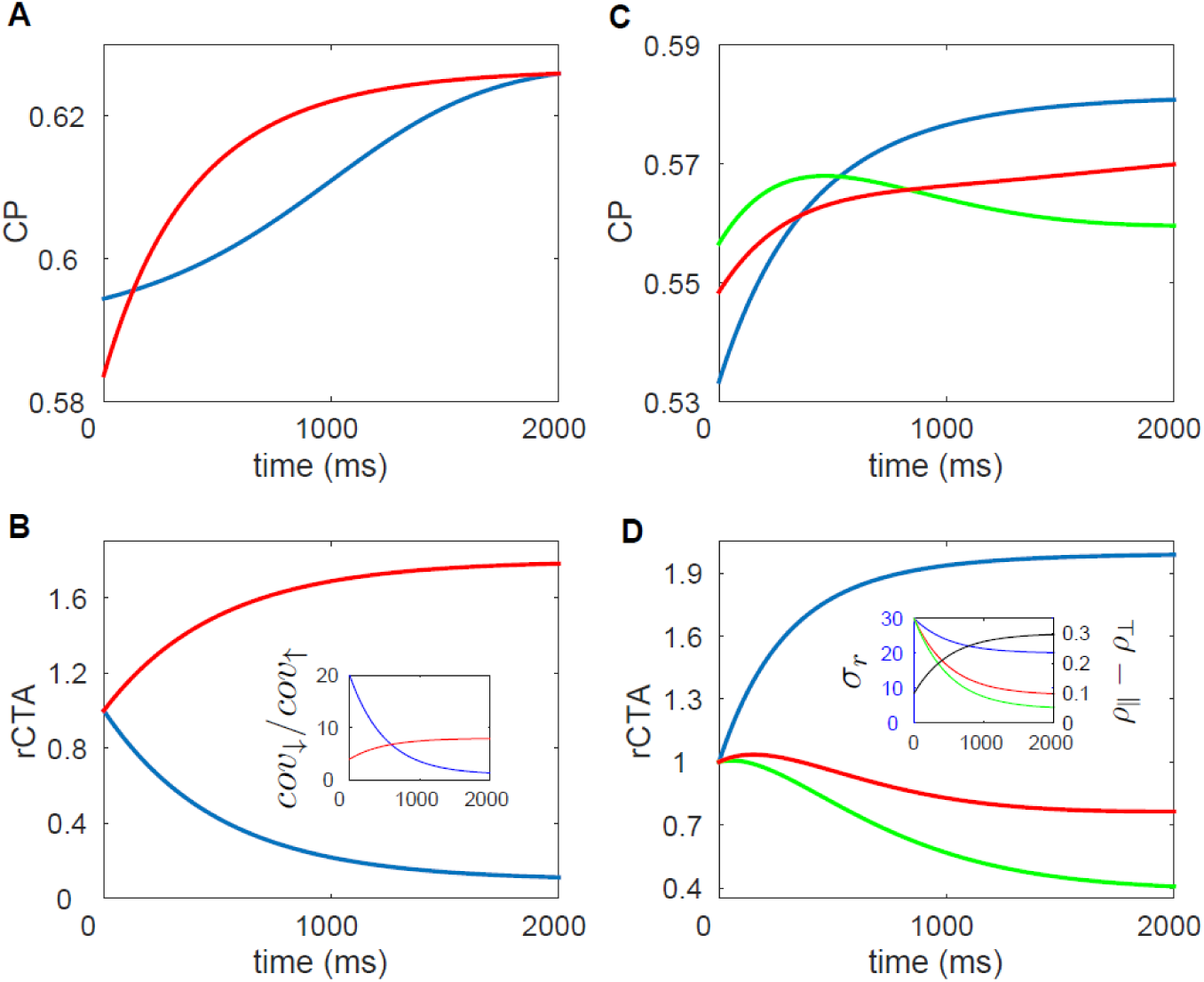
Temporal changes in the strength of different sources of activity-choice covariation affect differently the temporal profile of CPs and CTAs. A) CP temporal changes produced by the decrease in the influence of an internal variable with time (blue) and, alternatively, by the increase of the influence of decision-related feedback with time (red). See the main text for a rationale of these time dependencies for each of the two sources of covariation. B) Same as A) but for the relative CTA changes (rCTA), that is, the ratio of the CTA with respect to its value at time zero. Inset: Temporal profile of the ratio of the top-down (or feedback) contribution to the covariance cov(*r_i_*, d) and the feedforward contribution. C) CP temporal changes for a neuron-antineuron feedforward model produced by variations in the responses variance and in the difference of correlations between same or different types of cells, *ρ_||_ − ρ_⊥_.* D) Same as C) for rCTA. Inset: Temporal profile of the changes in the correlations difference (black line) and in response variance (coloured lines).

Fig. 6A shows that these changes in the weights have qualitatively similar effects on the CP, which increases over time in both cases. On the other hand, the effect on the CTA is the opposite (Fig. 6B). This can be understood considering the dependencies of cov(*r_i_*, *d*) and var *r_i_* separately (see Eqs. 29 and 30). The top-down and decision-related feedback contribution to *cov*(*r_i_, d*) that change with the weights, are (w^T^u + *w_a_*)*u_i_* and 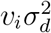, respectively. In the inset of panel B we show the temporal changes in the ratio between these contributions and the feedforward contribution to the covariance, which is assumed to be constant across time. CTAs change according to the changes in the covariance, but the CPs temporal profile depends also on the changes in var *r_i_*, given by 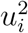 and 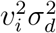, respectively. The CP increases in both cases because, in proportion to the response variability, cov(*r_i_*, *d*) increases. This occurs also when cov(*r_i_*, *d*) decreases with *u_i_* because the decrease in the influence of the internal variable also produces a decrease in the variability of the responses.

As a second example, we consider a feedforward model with a neuron-antineuron configuration (Shadlen et al., 1996; Cohen and Newsome, 2009; Nienborg and Cumming, 2010). This model contains two types of neuron: a set of neurons tuned to be more responsive to stimuli leading to *D =* 1, and a set of antineurons with the opposite tuning function, symmetrical with respect to the uninformative stimulus, and thus related to *d* = −1. Shadlen et al. (1996) showed by means of simulations that the CP in this case is mainly unsensitive to the distribution of the feedforward weights as long as they have opposite sign for the two sets. Furthermore, the CP is largely unsensitive to the responses variance, and is determined by the difference between the average noise correlations between neurons within the same set (*ρ*_||_) and noise correlations between neurons of different types (*ρ_⊥_*)(Nienborg and Cumming, 2010). These results were analytically confirmed by Haefner et al. (2013). We here examined time-resolved activity-choice covariations for this purely feedforward neuron-antineuron model (see Methods for details).

Fig. 6C-D show the CP and CTA for three cases in which the difference between the correlations (*ρ*_||_ − *ρ*_⊥_) increases with time and is accompanied by a decrease in the responses variance. These trends mimic the plausible scenario in which after stimulus onset there is a transient response with higher rates followed by some decrease of the activity during a tonic phase. For Poisson-like responses this leads to a decrease in the variance, which is proportional to the mean rate. On the other hand, the increase of *ρ*_||_ − *ρ*_⊥_ can occur as a consequence of recurrent connections that amplify the initial differences because their structure is related to tuning functions' similarity. The three cases shown in Fig. 6C-D share the same change in *ρ*_||_ − *ρ*_⊥_, while the variance changes are different (see inset panel). We see that, once the window used to estimate the responses does not cover the whole trial, the CP is not anymore independent of the responses variance, as had previously been found for this model. This is because when the CP is estimated only for a certain interval it matters which is the portion of the overall variance of the responses related to that interval (See Eq.16 in Methods for details). Regarding the comparison of time profiles for CPs and CTAs, we see that the combination of the same change in *ρ*_||_ − *ρ*_⊥_ with different changes in the response variance leads to different combinations of increasing or decreasing CPs and CTAs, reflecting that the measures characterize complementary aspects of the changes in the acticity-choice covariations.

Overall, these examples illustrate that to characterize the changes in activity-choice covariations it is useful to jointly study other measurable quantities complementary to the CP. Nienborg and Cumming (2009) fruitfully combined CPs with psychophysical kernels, which quantify the stimulus information impact on the choice. We here considered the CTA and the response variance because of their relation to the CP in its linear approximation. Moreover, jointly measuring the changes in response autocorrelations and, for simultaneous recordings, in noise correlations, can further characterize the changes in activity-choice covariations.

Furthermore, although in general the decision variable is not directly measurable, it may be estimated with novel experimental designs that allow controlling more precisely the amount of sensory evidence by discretizing its presentation (Brunton et al., 2013). Pitkow et al. (2015) proposed to estimate the variance of the decision variable indirectly, calculating the psychophysical threshold from the choice ratio (Eq.25) and assuming unbiased decoding. Estimating this variance is especially relevant to characterize the changes in activity-choice covariations when, in contrast to temporal changes examined here, changes across conditions are studied. Different conditions can be associated with different reward levels (Nienborg and Cumming, 2009), or with the inactivation of some areas (Smolyanskaya et al., 2015; Katz et al., 2016). In that case, the CTA changes do not only depend on changes in cov(*r_i_*, *d*) but also on changes in the variance of the decision variable.

#### 4.4 The characterization of feedback signals

Given the CTAs solutions we have obtained for the feedback models, in this section we show how stimulus dependencies of activity-choice covariations and their relation to tuning properties may help to distinguish between different theories about the nature of the feedback signals. To set a classification of different types of activity-choice covariations we consider the dependence of the sign of the CTA on the cells tuning properties. The choice preferences of a cell are determined by its relative responsiveness to stimuli informative about each choice. For example, a sensory neuron whose tuning function around the decision boundary stimulus (*s* = 0) has a positive slope (Fig. 7A) is expected to contribute to the decision variable with a positive readout weight and higher activity in the neuron will be indicative of choice *D* = 1 being made. Given the cells choice preferences, we ask if the feedback sign depends on these preferences and, if it does, wether feedback enhances or suppresses the responses of cells with tuning properties affine to each choice.

**Figure 7:**
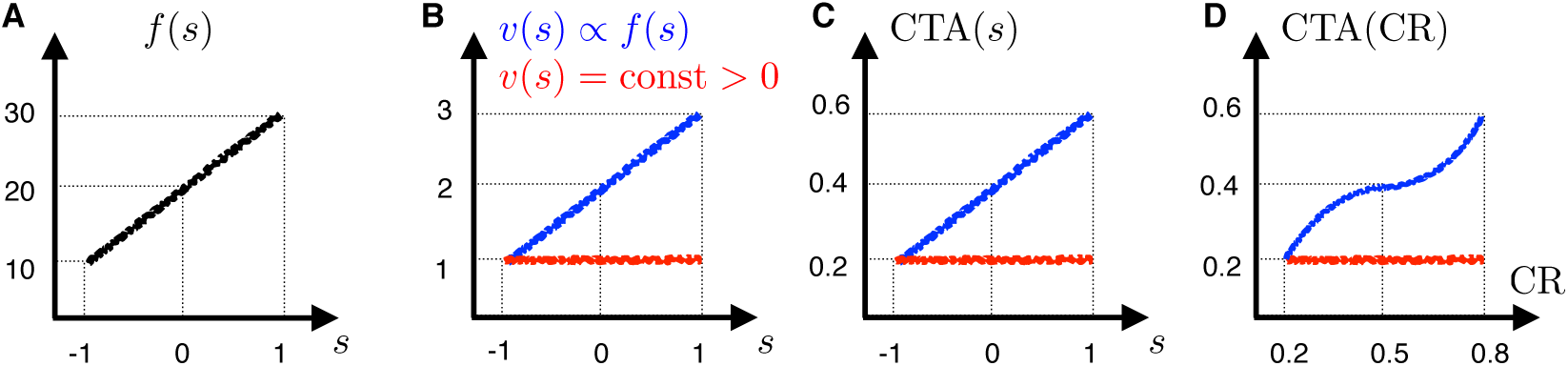
Illustration of how different types of FB signals shape the CTA dependence on the stimulus. **A)** Cartoon of an unmodulated feedforward tuning function around a zero-signal stimulus. **B-D**) Excitatory feedback models with an additive constant (red) or multiplicative (blue) influence on the sensory response, respectively. **B**) Feedback weight dependence on the stimulus. **C**) CTA dependence on the stimulus implied by B). **D**) Corresponding CTA dependence on the choice ratio (CR).

Based on this criterion, we can for example distinguish between predictive coding theories (Rao and Ballard, 1999) and probabilistic inference theories (Lee and Mumford, 2003; Fiser et al., 2010; Nienborg and Roelfsema, 2015; Haefner et al., 2016) of feedback. In predictive coding the feedback signals represent predictions at a higher level that are subtracted from the feedforward ones, so that sensory activity communicates a prediction error. Accordingly, the feedback is always negative, independently of the cell tuning. However, the sign of the CTA depends on the cell choice preferences. For a stimulus *s* > 0, cells with a positive tuning function slope, compatible with *D* = 1 as in Fig. 7A, are expected to receive on average a stronger negative feedback when choice *D* = 1 is made, because the prediction error is then smaller. Therefore, these cells will have CTA < 0, in contrast to cells with a reversed slope.

On the other hand, if sensory responses represent posterior beliefs, feedback signals represent prior information about the stimulus leading to an enhancement of neural activity compatible with the prior. In this case, the sign of the feedback for each cell may depend on the specific form of the prior. If the prior does not represent the details of the stimulus value but only task relevant information for the choice (s > 0 or not), then the feedback is categorical in nature and cells compatible with *D* = 1 as in Fig. 7A would receive a positive feedback for *D* = 1 and a negative one for *D* = − 1, leading to CTA > 0, and oppositely for cells with reversed slope. If the prior more locally determines the expected stimuli, activity may be enhanced also only for cells with more specific tuning properties and not only according to their choice preferences. However, also in this latter case, the relative average strength of the feedback for the two choices is expected to result in CTA > 0 for cells with choice preferences as in Fig. 7A. Altogether, the relation between the sign of the CTA and the cell choice preferences provides a signature to distinguish the effect of predictive and belief-related feedback signals.

Furthermore, the dependence of the CTA on the stimulus may further discriminate between different types of feedback. For example, probabilistic inference theories differ in how neural responses represent probabilities. It has been suggested that sensory responses represent posterior beliefs either as probabilities (Anastasio et al., 2000; Buesing et al., 2011) or as logarithms of probabilities (Ma et al., 2006; Jazayeri and Movshon, 2006), so that the feedforward (likelihood) and feedback (prior) signals would be combined multiplicatively or additively, respectively. The nature of the posteriors computation may be distinguished from the CTAs. In particular, consider the case mentioned above in which the belief is mostly about the choice, resulting in a categorical feedback. The CTA will mainly reflect the difference between the prior values for *D =* ±1, in agreement with *v_i_* ∆_*D*_ in Eq.31. If the prior is additively combined, the feedback does not depend on the stimulus (Fig. 7B). Oppositely, if the prior is combined multiplicatively, its strength is proportional to the feedforward tuning function. These dependencies are reflected in the CTAs (Fig. 7C-D). Therefore, for a categorical belief-related feedback we can identify how probabilities are represented studying the stimulus dependencies of the CTAs. Note however that, while this type of beliefs may be related to serial dependencies between choices made in subsequent trials, beliefs about the actual value of the sensory stimulus are expected to play a role during the integration of sensory evidence. This suggests that responses in early trial times may reflect more transparently the relation between CTAs and tuning properties characteristic of alternative probability representations.

#### 4.5 A concrete example of CTA analysis: Discrimination between a decision-related attention feedback model and a feedforward model with optimal readouts

We here study and compare two concrete models, namely a feedback model based on decision-related attention and a feedforward model with optimal readout weights. Using as an example a two-alternative forced choice orientation task, we derive the CTAs and examine their dependence as a function of the sensory stimulus. Furthermore, we consider the relation between the CTAs and the tuning functions. We show how the population CTA, i.e. the set of the CTAs arranged in relation to the tuning functions, characterizes the nature of the feedback. We already discussed above the different dependence for predictive coding or probabilistic inference between the CTAs and the cells choice preferences. Generally, if the decision-related feedback has a functional role, it is expected that its distribution across the population is related to the tuning similarity of the neurons. In particular, when the stimulus space is mapped to the neural responses so that neurons can be labeled according to their preferred stimulus, then CTAs can be studied jointly as a function of the presented stimulus and of the preferred stimulus that characterizes the responses of each cell. The comparison of the two concrete models here analyzed will allow us to show how the population CTA lies, in the space of the population responses, in the direction determined by the changes in the mean responses induced by decision-related responses components. Furthermore, we will indicate specific alternative predictions from the models which can be used experimentally to discriminate between these different sources of activity-choice covariation.

We start examining the feedback model. This model represents attention (Maunsell and Treue, 2006) by introducing a multiplicative attention-related gain that modulates the tuning curves. We further assume that attention is decision-related so that, once the choice has been made, the attended stimulus *s_att_* is the estimated stimulus associated with the decision variable, *ŝ(d)*. This link between *s_att_* and *d* introduces the decision-related feedback. Furthermore, we assume the estimation to be unbiased, so that on average the attended stimulus matches the presented stimulus (〈*s_att_*〉 = 〈*ŝd*〉 = *s*). The unmodulated feedforward mean responses of cell *i* depend on *s* − *s_i_* where *s_i_* is the preferred stimulus for which cell *i* is mostly responsive. On the other hand, the modulatory gain depends on *s_att_ − s_i_.* Accordingly, 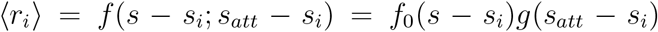. In terms of the general model of Eq.28, here *s*′ = *s_att_, f_i−FF_ =* f_0_(⋅), and *f_i−FB_ =* f_0_(⋅)[g(⋅) − 1], so that the feedback acts as a multiplicative gain.

For a concrete model implementation we chose an orientation task in which a visual stimulus with an angle *φ* is presented and the animal is asked to indicate on each trial if this angle is lower or higher than a boundary angle, which we take to be *π* (Fig. 8A). Depending on how informative the presented stimulus *φ* is, i.e. depending on |*φ* − π|, the choice ratio changes jointly with the average predicted angle, and thus with the average attended angle. We modeled the tuning curves with Von-Mises functions (Amirikian and Georgopulos, 2000; Ecker et al., 2016) and derived the CTAs using the linear approximation of Eq.28 (see Methods for details). In Fig. 8B we show the unmodulated and attention-modulated mean population response, with cells labeled according to their preferred stimulus (*φ_i_*). We also show the population CTA, that is, the CTA as a function of the preferred angle *φ_i_.* The CTA of each cell depends on the presented stimulus and on the preferred stimulus of the cell. In particular:

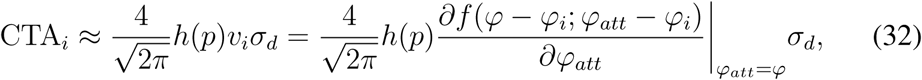

following the general expression of the CTA for continuous decision-related feedback (Eq.30) considering the form of the feedback weights for neural responses of the type described in Eq.28, namely proportional to the derivative *(f′_att_)* of the tuning function with respect to *φ_att_.* The association between the estimated and attended angles introduces the activity-choice covariations.

**Figure 8:**
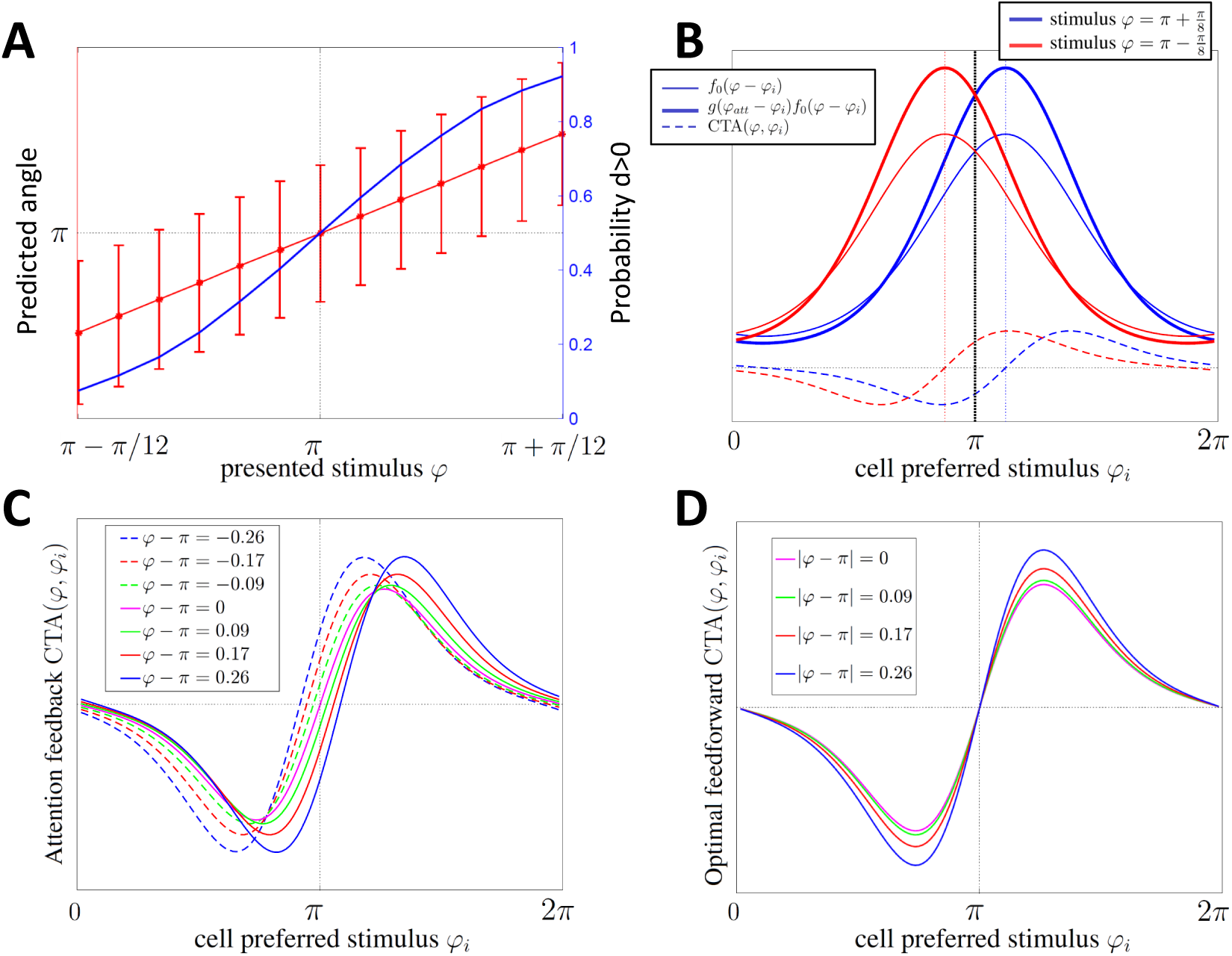
Comparison of activity-choice covariations between alternative feedforward and feedback models based on population CTA analysis for a representative two-alternative forced choice orientation task. **A**) Predicted angle (red line) and choice ratio (blue line) as a function of the angle *φ* of the stimulus presented. The predicted angle is estimated from the continuous decision variable d, depending on the stimulus information content. The black dotted line indicates the discrimination boundary (the uninformative stimulus *φ* = π). **B**) Post-decision population responses in the presence of decision-related attention-based feedback. Cell responses are shown as a function of the preferred stimulus of each cell (*φ_i_*) for two presented sensory stimuli. In particular, the unmodulated mean population responses, gain-modulated mean population responses, and CTAs are shown for *φ* = π ± π/8. Colored vertical dotted lines indicate the population response peak for each sensory stimulus. See Eq.18 in Methods for details on the responses model. **C)** Population CTAs in the presence of decision-related attention-based feedback. CTAs are displayed as a function of the preferred stimulus of each cell for a set of different sensory stimuli. **D**) Same as C) but for a feedforward model with optimal read-out weights.

The population CTA is examined in more detail in Fig. 8C for different presented stimuli. It changes with the presented stimulus in three ways: First, since the population response peak shifts with the presented stimulus, also the population CTA is shifted, being *f′_att_* = 0 at the peak (Fig. 8B). Second, the derivative *f*′_*att*_, and hence the CTA, has the same magnitude and opposite sign for angles symmetrically equidistant from the mean attended angle, corresponding to *φ*. That is, *f′_a_*_tt_(*φ* − *x*) *= −f′_a_*_tt_ (*φ* + *x*) (see Eq.21 in Methods for details). Accordingly, the population CTA reverses its sign and is mirrored around *φ_i_* = *φ*, for which CTA_*i*_ = 0. Third, the factor *h*(*p*) introduces a certain gain to the population CTA which is common to all cells and exclusively depends on the choice ratio as determined by | *φ* − *π* |.

We now compare the CTAs of this feedback model with the ones of a feedforward model with an optimal read-out. Haefner et al. (2013) determined the form of the CP when the feedforward weights are optimal for a two-choice discrimination task and derived an optimality test that was then applied by Pitkow et al. (2015) explicitly considering an optimal continuous estimator of the sensory stimulus. For this latter case, the optimal weights are (Pitkow et al., 2015): w ∝ C^−1^*f*′_*S*_, where *f′_s_* refers to the tuning function derivative with respect to the presented stimulus. Introducing these weights in the CTA expression for a feedforward model (Eq.27) in the orientation task the CTAs are

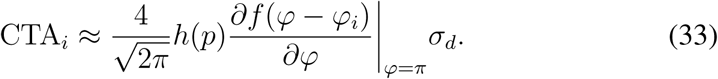

These CTAs only differ from the ones of the feedback model (Eq.32) in the derivative f′_s_ that replaces *f′_att_.* For an optimal decoder the population CTA is proportional to f′_s_ because this is the direction encoding changes in the presented stimulus and hence the direction providing optimal information for the choice. In both cases the population CTA lies in the direction of the changes in mean responses induced by decision-related responses components. In the optimal readout feedforward model these components are driven by the presented stimulus, while in the attention feedback model they are driven by the attended stimulus.

In Fig. 8D we show the population CTA obtained for the optimal read-out feedforward model when different stimuli are presented. For *φ* = π the population CTAs of Fig. 8C-D coincide because the multiplicative nature of the feedback in the attention model and the specific form of the feedforward and feedback components of the tuning functions lead to *f*′_*s*_ and *f*′_*att*_ having the same shape. However, for stimuli *φ* ≠ π, the population CTAs differ between the two models and for the optimal read-out feedforward model the presented stimulus only modulates the CTA through the factor *h*(p), with the choice ratio p determined by |*φ* − π|. The CTA is zero always for the cells which preferred stimulus is at the boundary (*φ_i_* = π), independently of the stimulus presented. These differences provide experimentally testable predictions to discriminate between these two alternative sources of activity-choice covariation.

### 5 Discussion

We derived a novel and general analytical expression of measures of covariation between neural activity and behavioural choice that is valid regardless of the source of the covariation for two-alternative choice tasks, comprising discrimination and detection tasks. Our framework extends previous analytical results assuming purely feedforward (FF) sources of this covariation by considering feedback (FB) sources of covariation as well as informative stimuli. We related several measures of activity-choice covariation including choice probability (CP) (Britten et al., 1996), choice correlation (Pitkow et al., 2015), and choice triggered average (CTA) (Haefner, 2015). The advantages of this analytical framework include a deeper understanding of the effect of various possible sources of dependence between the neural activity and behavioural choice, the possibility to formulate experimentally testable hypotheses about the form of the covariations for different sources -for example about their dependence on the sensory stimulus and their relation with the neural tuning properties- and to better understand what each statistical measure may tell us about the cause and changes in activity-choice covariations. Furthermore, our results substantially expand the applicability of the analytical relationships from discrimination to detection task, from zero-signal to non-zero signal trials, and from purely FF scenarios to those in which FB signals are important.

### Including the effect of sensory signals in activity-choice covariation analysis

A first advance that we made was to derive how activity-choice covariations are affected by task-informative signals contained in the sensory stimulus, independently of the origin of those covariations. We derived an exact CP solution under the assumption of Gaussian response variability and a more general linear approximation to the CP in terms of the CTA. Both formulas account for the effect of informative signals or choice biases that modify the ratio of selected choices. When the choice is mediated by an internal estimator of the sensory stimulus (Gold and Shadlen, 2007), we found that these measures depend on the stimulus signal always through a single multiplicative factor that only depends on the choice ratio. This factor is generic, in contrast to additional source-specific dependencies on the stimulus signal that may affect the covariations, for example if noise correlations are stimulus-dependent.

Previous work on how to combine neural responses to stimuli with different information content concentrated on discounting the effect of stimulus tuning on neural activity to estimate a genuine choice-related signal, for example either z-scoring the responses (Kang and Maunsell, 2012) or subtracting out their estimated stimulus-driven component (Nienborg and Cumming, 2009), assuming that this contribution is additive. Pooling data from all stimulus conditions should allow a more reliable estimation of CPs (e.g. Padoa-Schioppa, 2013; Pitkow et al., 2015). Here we characterized the generic multiplicative factor *h*(*p*) and verified that its Gaussian approximation is robust if a large population of neurons contributes to the decision. Accordingly, we provided an alternative way to factor out this influence and combine data from different signal levels with minimal assumptions about the origin of the activity-choice covariation. Furthermore, our work suggests that an even better way to make full use of the data is to study CPs and CTAs as a function of the stimulus, since the form of this dependence helps disambiguating between different sources.

### Beyond feedforward models of choice: how the form of activity-choice covariations reflects different sources of choice-related activity

The second advance we made was to compute CPs and CTAs under general conditions that include feedback sources of activity-choice covariation. We examined different causal architectures of the connections between neural activity and choice and characterized the effect of feedback signals either directly related to the decision process or to other internal variables, such as slow fluctuations of neural excitability (Ecker et al., 2014), attention (Maunsell and Treue, 2006), and prediction (Rao and Ballard, 1999) or belief-related (Lee and Mumford, 2003; Haefner et al., 2016; Tajima et al., 2016) signals. These non stimulus-driven signals can lead to across-trials dependencies in activity and behavior (Frund et al., 2014; Conen and Padoa-Schioppa, 2015; Nienborg and Macke, 2014) and cause anticipatory activity-choice covariations before the stimulus is presented (Padoa-Schioppa, 2013).

Our CP and CTA solutions allow identifying different qualitative ways in which internal states can contribute to activity-choice covariations. First, feedback signals may affect noise correlations (Cohen and Maunsell, 2009; Mitchell et al., 2009; Wimmer et al., 2015; Ecker et al., 2016; Haefner et al., 2016) and propagate onto the decision through the feedforward readout, hence modifying feedforward contributions to activity-choice covariations (Cumming and Nienborg, 2016). Second, internal states may commonly drive neural activity and the choice. For decision-related feedback, the decision variability itself modulates the responses. A common signature of activity-choice covariations produced by feedback signals is that, oppositely to feedforward covariations which rely on noise correlations of a certain cell with the rest of the population, significant covariations may arise from the direct effect of the feedback signal on that cell. In particular, if decision-related feedback is dominant, CTAs proportionally reflect the strength of the feedback weights.

The characteristic form of CPs and CTAs may help to discriminate between alternative hypotheses regarding the source of activity-choice covariations. In a concrete example, we showed how to use CTAs to distinguish between a feedback model with decision-related attention and a feedforward model with optimal readout weights. These two models result in different population CTA dependencies as a function of the presented stimulus, which can be tested experimentally. More generally, we also discussed how to characterize from CTAs whether the primary effect of feedback is multiplicative or additive, and whether enhances or supresses neural activity, to gain further insights on what is the function of the feedback signals, for example whether carries error signals (Rao and Ballard, 1999), or represents probabilities (Buesing et al., 2011; Haefner et al., 2016) or log-probabilities (Ma et al., 2006; Jazayeri and Movshon, 2006; Tajima et al., 2016) in probabilistic codes.

### The complementary information of measuring CTAs as well as CPs

Since Britten et al. (1996), CPs have been widely used to quantify activity-choice covariations, in contrast to CTAs. While CP is sensitive to higher orders of divergence between the distributions of responses for each choice, the CTA captures the linear differences. We showed that this reduced, more specific, sensitivity confers the CTA with desired properties such as additivity and linearity, resulting in simpler dependencies useful to infer feedback weights and allowing to derive tractable expressions for hypothesized models. Only in CTAs feedback and feedforward contributions combine additively, allowing for a simpler disentangling of their relative contributions. The CPs capture both the effect of response variance and of covariations between the responses and the choice, whereas CTAs captures only the latter. The complementarity of measuring both CPs and CTAs is particularly appealing for examining the time changes in activity-choice covariations (Nienborg and Cumming, 2009). As illustrated in our examples, teasing apart changes in the response variance and in the covariance between the responses and the choice helps to distinguish between effects like the decay of serial dependencies in the late part of the trial, the increase of feedback influences in the late part, and changes in response variability associated with transient responses after the stimulus onset.

Our analytical approach also identifies other factors that can often be measured and can further help to pin down the exact sources of activity-choice covariations. These factors include noise correlations (Smolyanskaya et al., 2015), autocorrelations, and the amount of trial-to-trial variability in the decision variable, which can be estimated indirectly from the psychophysical threshold (Pitkow et al., 2015).

### Taking advantage of stimulus manipulations

The inclusion of informative signals and feedback in our formulation suggests how experimental designs exploiting stimulus manipulations can further improve the discrimination of activity-choice covariation sources. One prediction from the comparison between continuous and categorical feedback models is that only in the presence of continuous feedback the CTA is sensitive to the manipulation of the stimulus trial-to-trial variability. Furthermore, given the generality of our derivations, results can easily be extended to compare response distributions associated with a certain range of reaction times or of values of the internal decision variable that integrates sensory evidence, thus allowing the assessment of confidence degrees (Kiani and Shadlen, 2009). Moreover, the manipulation of the stimulus duration and of the delay interval before the choice (e. g.Luna et al., 2005; Kiani et al., 2008) can modulate the relative influences of feedforward and feedback sources in different parts of the trial, making it possible in principle to use our formulas to infer feedforward and feedback weights independent of each other, constraining models of cortical computations.

## Acknowledgments

This work was supported by the Fondation Bertarelli and by the Autonomous Province of Trento, Call Grandi Progetti 2012, project Characterizing and improving brain mechanisms of attention-ATTEND.

## Footnotes

1 This relation holds for the covariance between any variable *x* and a binary variable *D*, and independently of the convention adopted for the values of *D:* the factor 2 has to be replaced by *a – b* if *D* = *a*, *b* is selected instead of *D* = 1, –1.

